# Human iPSC-derived neuron of 16p11.2 deletion reveals haplotype-specific expression of *MAPK3* and its contribution to variable NDD phenotypes

**DOI:** 10.1101/2022.07.10.498576

**Authors:** Fang Liu, Chen Liang, Zhengchang Li, Sen Zhao, Shaofang Shangguan, Haiming Yuan, Ruen Yao, Zailong Qin, Shujie Zhang, Liping Zou, Zhijie Gao, Qian Chen, Suiwen Wen, Jing Peng, Fei Yin, Fei Chen, Xiaoxia Qiu, Jinsi Luo, Yingjun Xie, Dian Lu, Yu Zhang, Hua Xie, Hongying Wang, Xiaodai Cui, Jian Wang, Hailiang Huang, Ruize Liu, Xiaofang Sun, Chao Chen, Nan Wu, Chunyu Liu, Yiping Shen, James F. Gusella, Xiaoli Chen

**Affiliations:** Department of Medical Genetics, Capital Institute of Pediatrics, Beijing 100020, China; Graduate School of Peking Union Medical College, Beijing 100730, China; the Department of Orthopedic Surgery, Key Laboratory of Big Data for Spinal Deformities, Beijing Key Laboratory for Genetic Research of Skeletal Deformity, State Key Laboratory of Complex Severe and Rare Diseases, all at Peking Union Medical College Hospital, Peking Union Medical College and Chinese Academy of Medical Sciences, Beijing 100730, China; Dongguan Maternal and Child Health Care Hospital and Dongguan Institute of Reproductive and Genetic Research, Dongguan 523120, China; Department of Medical Genetics, Shanghai Children’s Medical Center, Shanghai Jiao Tong University School of Medicine, Shanghai 200127, China; Genetic and Metabolic Central Laboratory, Birth Defect Prevention Research Institute, Maternal and Child Health Hospital of Guangxi Zhuang Autonomous Region, Nanning 530002, China; Department of Pediatrics, Chinese PLA General Hospital, Beijing 100039, China; Department of Neurology, the affiliated hospital of Capital Institute of Pediatrics, Beijing 100020, China; Department of Obstetrics, Qingyuan People’s Hospital, 6th Hospital of Guangzhou Medical University, Qingyuan 511518, China; Department of Pediatrics, Xiangya Hospital of Central South University, Hunan Intellectual and Developmental Disabilities Research Center, Changsha 410008, China; Key Lab for Major Obstetric Diseases of Guangdong Province, Experimental Department of Institute of Gynecology and Obstetrics, Third Affiliated Hospital of Guangzhou Medical University, Guangzhou 510150, China; Department of Lab center, Capital Institute of Pediatrics, Beijing 100020, China; Department of Clinical Laboratory, Children’s Hospital of Soochow University, Suzhou 215000, China; Analytic and Translational Genetics Unit, Massachusetts General Hospital, Broad Institute of MIT and Harvard, Cambridge MA 02142, USA; Center for Medical Genetics & Hunan Key Laboratory of Medical Genetics, School of Life Sciences, Central South University, Changsha 410083, China; Department of Psychiatry, SUNY Upstate Medical University, NY 13201, USA; Division of Genetics and Genomics, Boston Children’s Hospital, Department of Neurology, Harvard Medical School, Boston, MA 02115, USA; Molecular Neurogenetics Unit, Center for Genomic Medicine, Massachusetts General Hospital, Boston, MA 02114, USA; Blavatnik Institute, Department of Genetics, Harvard Medical School, Boston, MA 02115, USA

**Keywords:** 16p11.2del, human induced pluripotent stem cell (hiPSC), neurodevelopmental disorder, residual haplotype, enhancer SNP, MAPK3

## Abstract

Recurrent proximal 16p11.2 deletion (16p11.2del) is risk factor of diverse neurodevelopmental disorders (NDDs) with variable penetrance. Although previous human induced pluripotent stem cell (hiPSC) models of 16p11.2del confirmed disrupted neuron development, it is not known which gene(s) at this interval are mainly responsible for the abnormal cellular phenotypes and how the NDD penetrance is regulated. After haplotype phasing of 16p11.2 region, we generated hiPSCs for two 16p11.2del families with distinct residual haplotypes and variable NDD phenotypes. We also differentiated the hiPSCs to cortical neural cells and demonstrated *MAPK3* as a driver signal of 16p11.2 region contributing to the dysfunctions in multiple pathways related to neuron development, which leads to altered morphological or electrophysiological properties in neuron cells. Furthermore, residual haplotype-specific *MAPK3* expression was identified in 16p11.2del neuron cells, associating *MAPK3* down-expression with the minor allele of the residual haplotype. Ten SNPs of the residual haplotype are mapped as enhancer SNPs (enSNPs) of *MAPK3*, eight enSNPs were functionally validated by luciferase assays, implying enSNPs contribute to residual haplotype-specific *MAPK3* expression via cis-regulation. Finally, the analyses of three different patient cohorts showed that the residual haplotype of 16p11.2del is associated with variable NDD phenotypes.

## INTRODUCATION

Neurodevelopmental disorders (NDDs) are a group of conditions characterized by cognitive, neurological and/or psychiatric manifestations due to abnormal development of neural system. Among the known genetic etiologies for NDDs, recurrent 16p11.2 deletion at breakpoint 4 and 5 locus (chr16: 29.6-30.2Mb, abbreviated as 16p11.2del) has emerged as a risk factor. The phenotypes of 16p11.2del include autism spectrum disorder (ASD) (Weiss et al., 2008), intellectual disability/developmental delay (ID/DD) (Cooper et al., 2011; Moreno-De-Luca et al., 2015), Seizures (Shinawi et al., 2010), speech delays and behavioral problems (Niarchou et al., 2019; Rein and Yan, 2020) and psychosis (Steinberg et al., 2014).

Mouse models had been developed to study the effects of 16p11.2del on the central nervous system development and behavior (Portmann et al., 2014; Arbogast et al., 2016). Zebrafish and Drosophila models had also been developed to explore the driver genes of 16p11.2del for NDDs (Golzio et al., 2012; Iyer et al., 2018). Studies of animal models recapitulated some core behavioral phenotypes and identified some key biological pathways or molecular targets of 16p11.2del. For example, the ERK signaling, which has prominent roles in brain development, was altered in the deletion mice (Pucilowska et al., 2015), and the pharmacological inhibition of ERK signaling rescued the pathophysiology and abnormal behavioral phenotypes in deletion mice (Pucilowska et al., 2018). Knockout of *Taok2* resulted in cognitive and social deficits in mice with increased brain volume, as well as reduced dendritic growth and deficient excitatory synaptic transmission in prefrontal cortex (Richter et al., 2019). However, some phenotypes were inconsistent across models or opposite between mice and humans. For example, *kctd13* knockout affected the head size in zebrafish (Golzio et al., 2012) but not in the *Kctd13*-deficient mice (Escamilla et al., 2017) (Arbogast et al., 2019). The 16p11.2del mice exhibit reduced brain volume (Pucilowska et al., 2015), while deletion patients exhibited macrocephaly (Zufferey et al., 2012).

Human induced pluripotent stem cells (hiPSCs) are useful system for studying transcriptomics profiles, cell morphology, electrophysiological properties of mutant neuron cells. So far, hiPSC-derived neuron (Deshpande et al., 2017; Roth et al., 2020; Sundberg et al., 2021) of 16p11.2del have been created. Abnormal transcriptional alterations (Roth et al., 2020), increased soma size and dendrite length with reduced synaptic density were seen in 16p11.2del neurons (Deshpande et al., 2017). In addition, increased electrophysiological signatures were reported in 16p11.2del cortical neurons (Deshpande et al., 2017) and were replicated in dopaminergic neurons (Sundberg et al., 2021). These hiPSC models confirmed the direct effect of 16p11.2del to abnormal neurogenesis, yet, it is unknown which gene is the driver gene for abnormal cellular phenotypes.

Another unsolved issue of 16p11.2del is variable expression of NDD phenotype (Girirajan et al., 2012). Although previous study showed that the phenotypic variation of 16p11.2 can be partially explained by the presence of additional large CNVs (Girirajan et al., 2012; Duyzend et al., 2016), the contribution of deleterious mutation or regulator haplotype on the residual 16p11.2 copy to NDD heterogeneity has not been studied. We ascertained two 16p11.2del families with at least two carriers in each family (Shen et al., 2011; Xie et al., 2020). The intra-family carriers had discordant NDD phenotype. Although no significant coding variant was detected to explain the variable severity of NDD, different partial haplotype of residual 16p11.2 copy was revealed. With the advancement of ENCODE knowledge and genome-wide enhancer-related sequencing techniques, the regulation of non-coding variants to transcriptome and histone modification during neurogenesis have been revealed via 3D chromosome contact, transcriptional or regulator factors (Won et al., 2016; Girdhar et al., 2018; Song et al., 2019). Our 16p11.2del study regarding congenital scoliosis (CS) has demonstrated the residual haplotype defined by three *TBX6* SNPs (rs2289292T-rs3809624C-rs3809627A) added the risk of CS phenotype by hypomorphic effect (Wu et al., 2015). Two such non-coding SNPs locate on an elite regulatory element (GH16J030089) (Fishilevich et al., 2017), indicating their potential cis-regulatory roles. Therefore, characterizing the association between the residual haplotype and the expression of 16p11.2 genes, neuron development outcomes will decode the genetic mechanism of variable NDD phenotype observed in clinical.

In this study, we characterized the residual haplotypes for 31 NDD children with 16p11.2del, and generated hiPSCs and cortical neural cells for two 16p11.2del families showing intra-family distinct residual haplotypes and discordant NDD phenotypes. After analyzing transcriptomics, electrophysiological properties and morphologies of neuron cells at two developmental stages (neural progenitor cells and mature neuron cells), we identified *MAPK3* as the major gene of 16p11.2del contributing to disrupted transcriptomic profiles and neuron development. The residual haplotype specific-*MAPK3* expression and neuron phenotypes were observed in 16p11.2del neuron cells, showing lower *MAPK3* down-expression and worsen electrophysiological properties in neuron cell presenting minor residual haplotype. Further, we proposed and proved the hypothesis that the residual haplotype-specific expression of *MAPK3* is cis-regulated by ten SNPs on enhancer elements, which is associated with variable NDD phenotype of 16p11.2del.

## RESULTS

### The general information of Chinese child NDD cohort with 16p11.2del

This Chinese 16p11.2del NDD cohort of Capital Institute of Pediatrics (CIP-NDD-16p11.2del) consisted of 31 NDD who referred to Chromosomal microarray analysis CMA) in pediatric hospitals as previously reported (Yuan et al., 2021). There are 19 males (61%) and 12 females (39%) aged between 0.42-18 years. The inheritance of 18 child carriers (18/31=58%) was tested, nine deletions are inherited (maternal:4/31=13%; paternal:5/31=16%), and nine ones are *de novo* (9/31=29%) (Fig.1A).

**Figure 1.**
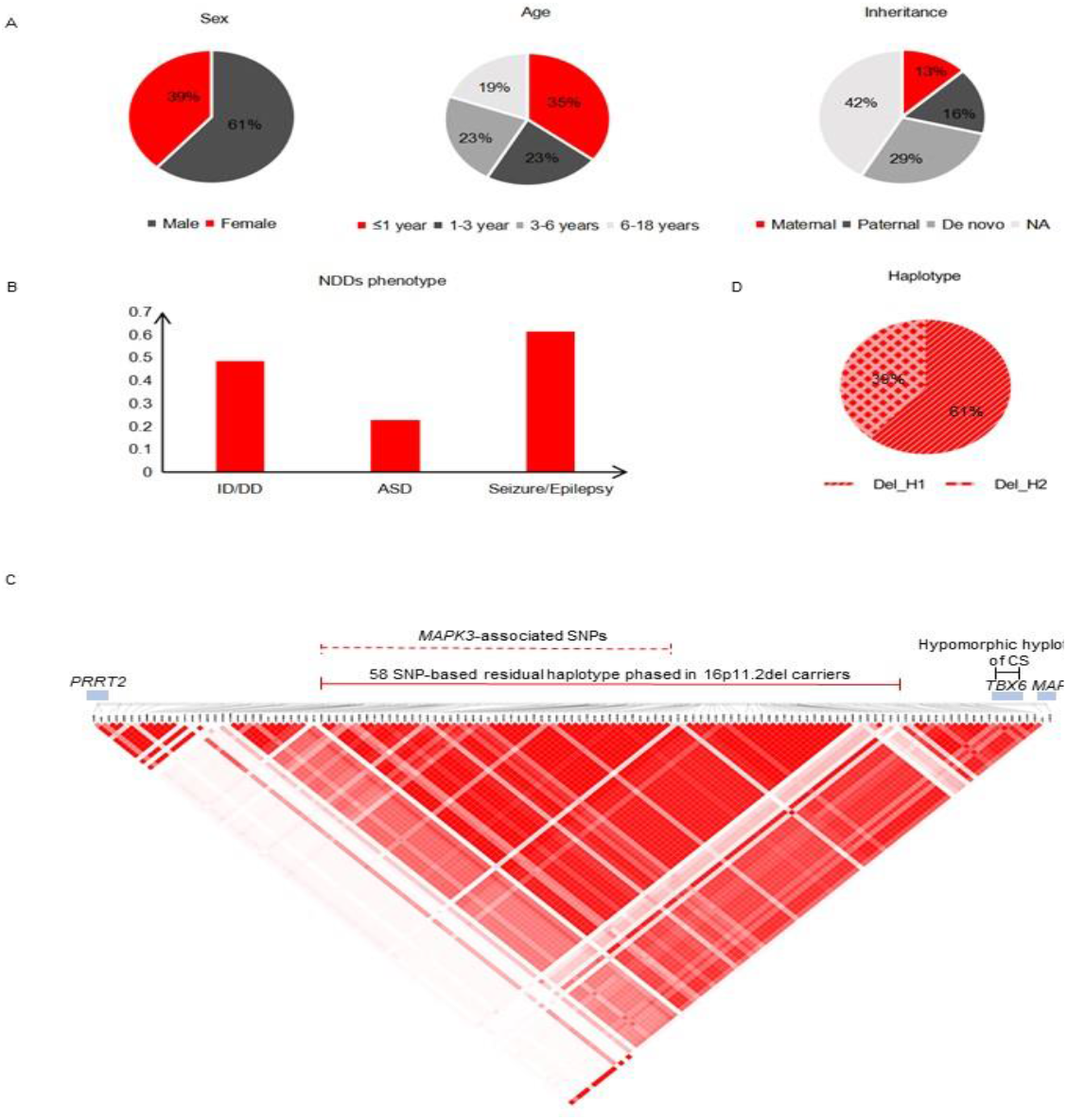
information of Chinese child NDD cohort with 16p11.2del. 31 NDD children with 16p11.2del from 29 Chinese families were summarized, showing the sex, age and inheritance (A), detailed NDD categories (B), linkage disequilibrium (LD) of 132kb 16p11.2 region (C) and frequency distribution of different 16p11.2 residual haplotypes(D). 58 SNPs consist the residual haplotype of 16p11.2 region. Del_H1: minor residual haplotype; Del_H2: major residual haplotype.

The NDD phenotypes of 31 child carriers is broad with much higher frequencies of developmental delay (48%) and seizure/epilepsy (61%) than ASD (23%) (Fig. 1B).

### Linkage disequilibrium (LD) of 132kb 16p11.2 region in the Chinese CIP-NDD-16p11.2del cohort

The residual haplotype at 16p11.2 locus was phased using captured sequencing (see Fig. 8A for probe coverage). 31 parents of these children (seven parents carrying 16p11.2del, 26 parents without 1611.2del) were available for capture sequencing to confirm the accuracy and Mendelian inheritance model of residual haplotype for NDD.

The SNPs of 16p11.2 region with MAF of 10-90% at homozygous status were extracted to phase the residual haplotype. Using online QCI software and Haploview software (Barrett et al., 2005), we identified 58 homozygous SNPs (Table S1) spanning 132kb (chr16:29924422-30057148) showed linkage disequilibrium in 31 children carriers and seven parental carriers (labeled as red solid line in Fig. 1C). As a result, we constructed two residual haplotypes in this cohort (Table 1). The reported CS-related *TBX6* hypomorphic alleles (chr16:30097630-30103160, rs2289292-rs3809624-rs3809627, labeled as black solid line in see Fig. 1C) (Wu et al., 2015) were behind this LD block. The MAF of the 58 SNPs ranged between 0.34-0.39 in 600 unrelated Chinese (Table 1). Interestingly, one residual haplotype consists of mutant/minor alleles (minor haplotype, abbreviated as Del_H1) while another one with wild-type/major alleles (major haplotype, abbreviated as Del_H2). The frequencies of Del_H1 and Del_H1 were 61% and 39% respectively in this cohort (Table 1 and Fig. 1D). In four families, the residual haplotypes of intra-family carriers are different, and such two families were chosen as iPSC donors.

**Table 1.**
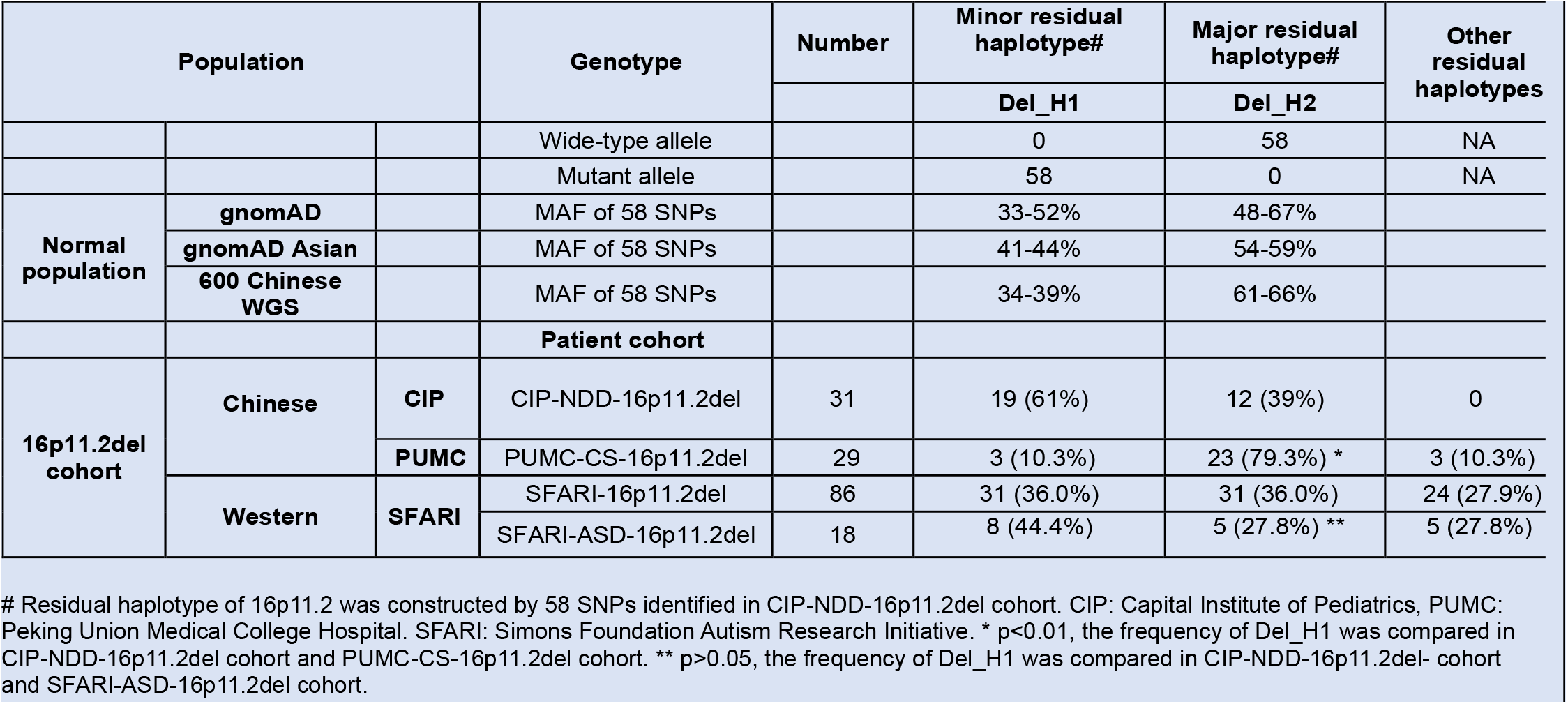
The frequencies of different residual haplotype in general populations and three patient cohorts with 16p11.2del.

We used the raw SNP chips of 216 inhouse Chinese and 160 European normal parents from Simons Foundation Autism Research Initiative (SFARI Base, https://www.sfari.org/) (Simons Vip, 2012) to construct the LD block of general population (Fig. S1). As showed in Fig. S1, 58 SNPs in the haplotype showed linkage disequilibrium in both Chinese and Western populations.

### Generation of human iPSCs from four 16p11.2del carriers and four controls

hiPSCs (Fig. 2A) were generated for six donors of two 16p11.2del families, including four deletion carriers (Del1, Del2, Del3, Del4), two normal parents (C2, C4). Besides, hiPSCs were generated also for two unrelated normal adults with different sexes (C1, C3), Two 16p11.2del families were chosen based on their unique advantages: I) The intra-family carriers are of same sex (mother/daughter pair in Family 1 and male sibs in Family 2) but the residual haplotypes of 16p11.2 are different. Del1 and Del4 are phased as Del_H1, while Del2 and Del3 as Del_H2. II) Carriers differ in severity of NDD features (Table S2) (Shen et al., 2011; Xie et al., 2020), which allow us to explore the relationships between the residual haplotypes and cellular phenotypes of neuron. III). All 16p11.2del donors don’t carry other rare genic CNVs >200kb, or deleterious single nucleotide variants according to the results of CMA and whole exome sequencing.

**Figure 2.**
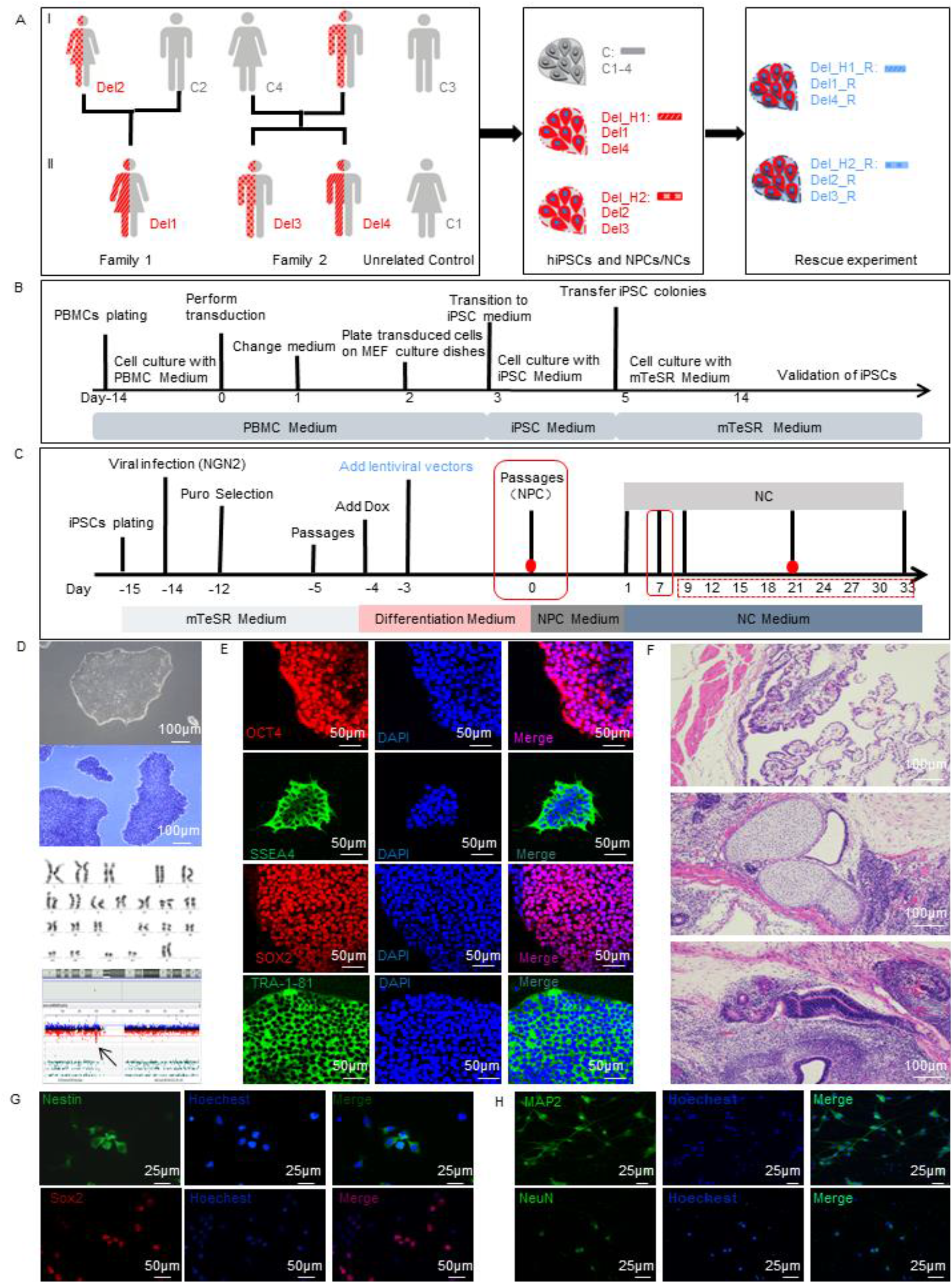
The experimental flowchart of hiPSC generation and cortical neural cell differentiation for16p11.2del carriers and controls. A, The pedigrees of two 16p11.2del families for hiPSC generation. Red symbol in body presented for 16p11.2del (abbreviated as Del) and grey symbol in body for control (abbreviated as C). Different symbols in body presented for different residual haplotypes in 16p11.2del, diagonal line for minor residual haplotype (Del_H1) while grid line for major residual haplotype (Del_H2). The hiPSCs were derived from PBMCs for six donors of two 16p11.2del families (Del1, Del2 and C1 in Family 1, Del3, Del4 and C4 in Family 2), and two unrelated controls (C1, C3). Father in Family 2 was not available for iPSC generation due to death in 51 years. The hiPSC-derived cortical neuron cell for MAPK3/PRRT2 rescue experiment were presented by blue colors (abbreviated as R). B, The timeline of hiPSC generation from PBMCs. C, The timeline of cortical neuron cell differentiation from hiPSCs and experiment. Day0: NPC stage after validation. The timepoints of RNA-Seq (Day0 and Day21), cellular morphology analysis (Day0 and Day7), electrophysiological recording (Day9-Day33) were labeled by red solid dots, red solid rectangles and red dashed rectangles. MAPK3/PRRT2 rescue experiment was performed at Day-3. D, Representative validation images of hiPSCs. White light of microscope (up panel). AP staining (middle panel). karyotype and the array-CGH analysis of hiPSCs (down panel). 16p11.2 deletion retained in hiPSCs (arrowhead). E, Images of hiPSCs markers OCT4, SOX2, SSEA4 and TRA-1-81. F, hiPSCs were differentiated into three germ layers: endoderm, mesoderm and ectoderm from top to bottom. The haematoxylin and eosin staining of teratomas derived from hiPSCs. G-H, Representative immunostaining images of NPC (Nestin and SOX2, G) and NC markers (MAP2 and NeuN, H) from derived neural cells.

PBMCs were reprogrammed to hiPSCs using standard protocol (Fig. 2B) (Seki et al., 2010). A total of 20 iPSC clones (Del: N=10; C: N=10) were successfully generated for eight donors with 2-3 clones for each donor (Table S2). The colonies have typical human embryonic stem cell morphology (Fig. 2D). All iPSC clones were examined for pluripotency markers (OCT4, SSEA4, SOX2 and TRA-1-81, Fig. 2E), and were validated for their differentiation capability to three germ lineages (Fig. 2F). Other quality control data (AP staining, karyotype and array CGH, Fig. 2D), mycoplasma testing and genomic identities (Table S3) were also performed for each iPSC clone.

### Differentiation of hiPSCs into cortical neuron cells

hiPSC colonies were differentiated into neurons with a previous protocol (Fig. 2C) (Zhang et al., 2013). The timepoint of successful differentiation to cortical neural progenitor cell (NPC) was defined as Day0, and the time before Day0 was labeled using minus Day. As shown in Fig. 2G-2H, positive staining of Nestin and SOX2, the markers of NPC, were detected on Day0 (Fig. 2G), while positive staining of the markers of neuron cell (NC) were detected on Day7 (MAP2) and Day21 (NeuN, Fig. 2H). The differential developmental profiles of NPCs and NCs were also demonstrated by the principal component analysis (PCA) of transcriptome (Fig. 3A). As expected, *NES* and *SOX2* expressed at a higher level in NPCs, whereas *DCX, MAP2* and *SLC17A6* expressed at *a* higher level in NCs (Fig. 3D). Sixteen transcriptomic data of previous hiPSC-derived 16p11.2del NPCs(Roth et al., 2020) (Fig. 3A) also clustered closely with our NPCs, but not with our NCs. Five of 16p11.2 genes (*C16orf92, TBX6, C16orf54, SPN* and *ZG16*) showed extremely low expression consistently in control hiPSC-derived neuron cells at both NPC and NC stage (FPKM<1), suggesting they unlikely to be relevant for early neurogenesis.

**Figure 3.**
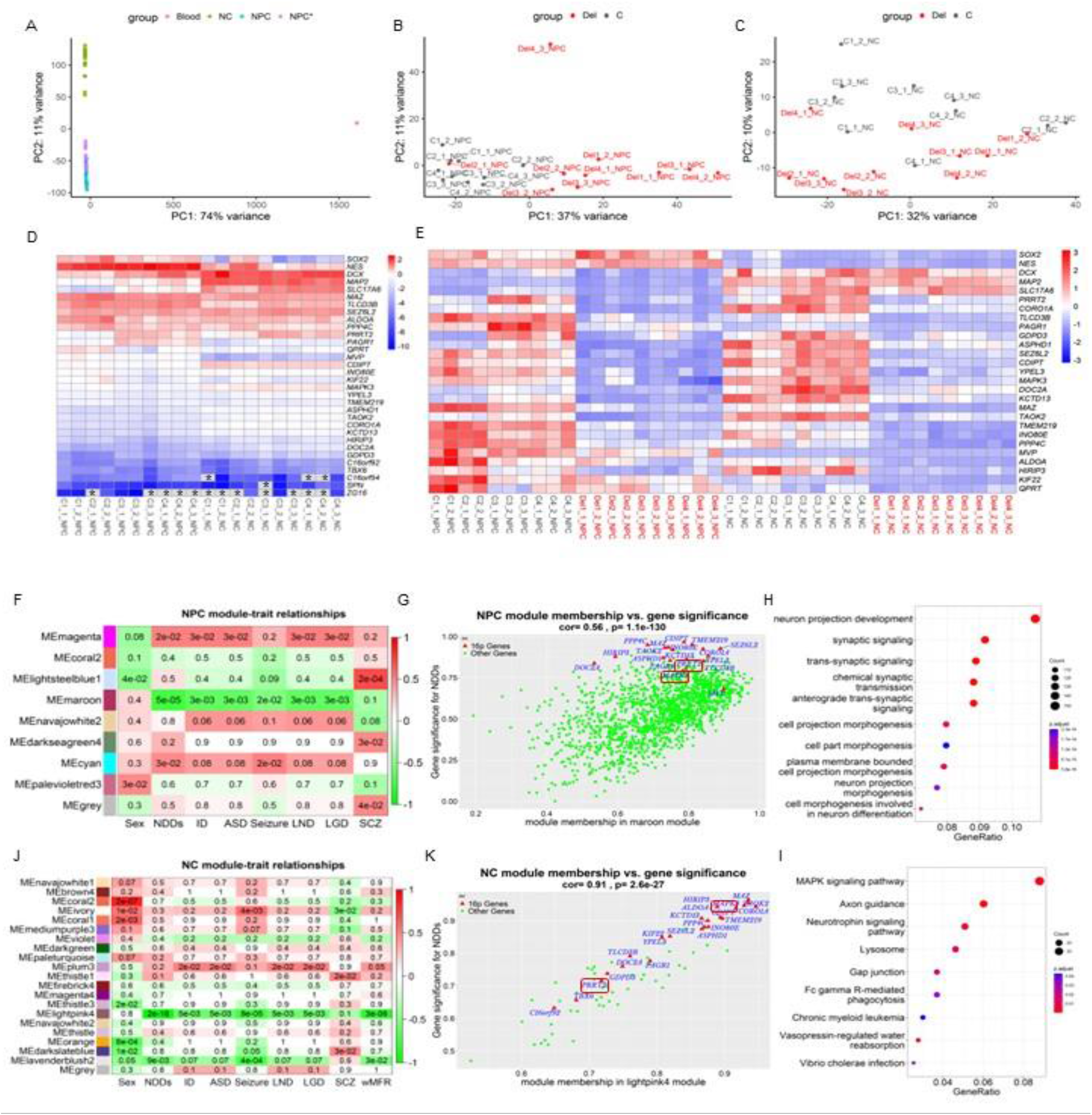
Transcriptomic profiles and WGCNA analysis of hiPSC-derived cortical neural cells with 16p11.2del. A-E, Time course transcriptome revealed differential neurogenesis of hiPSC-derived neural cells between NPC and NC stages, 16p11.2del carriers and controls. A-C, PCA of variance-stabilized count data after normalization and batch correction. Axes represent the first two principal components (PC1, PC2). A, Cell populations cluster by different cell type. Blood: N=1. NPC: N=20. NC: N=20. NPC*: NPCs from previous hiPSC study of 16p11.2del (Roth et al., 2020), N=16. B-C, Cell populations cluster by 16p11.2del status at NPC (B) and NC stages (C). Del: N=10, C: N=10. D, Heat maps of gene expression between normal NPCs and NCs, including NPC markers (NES and SOX2), NC markers (DCX, MAP2 and SLC17A6), and 16p11.2 genes. *Gene (FPKM=0) with extreme low expression in NPCs. E, Heat maps of gene expression between 16p11.2del carriers and controls, including neural cells markers and 16p11.2 genes. F-K, WGCNA revealed modules of co-expressed genes in hiPSC-derived cortical neural cells at NPC and NC stages. p<0.05 is statistically significant. F-G, Heatmap of p-values assessing the significance of module-trait correlations for NPCs. Module MEmaroon had the most significance and highest correlation with NDDs. LND, Learning difficulty. LGD, Language delay, SCZ, Schizophrenia(F). Visualization of gene significance (GS, y-axis), module membership (MM, x-axis) in MEmaroon (G). H-I, GO biological processes (H) and KEGG analysis pathway (I) enriched with genes belonging to MEmaroon of NPCs. J-K, Heatmap of p-values assessing the significance of module-trait correlations for NCs. Module MELightpink4 had the highest correlation with NDDs. LND, Learning difficulty. LGD, Language delay, SCZ, Schizophrenia (J). Visualization of gene significance (GS, y-axis), module membership (MM, x-axis) in the Module MELightpink4 (K). The gene list of Fig. 3H were in Table S11, the gene list of Fig. 3I were in Table S12.

### Dysfunctions in multiple pathways related to neuron development in cortical neuron cells with 16p11.2del

Transcriptomic comparisons were performed on two cell populations (Del, N=10, C, N=10) at two differentiation stages (NPC and NC stages). PCA was used to evaluate the quality of transcriptome with correction of experiment batch and additional uncharacterized factors. The expression of NPC markers (*NES* and *SOX2*) and NC markers (*DCX*, *MAP2* and *SLC17A6*) between 16p11.2delss and controls were similar (Fig. 3E and Table S4), confirming the success establishment of hiPSC-derived 16p11.2del neuron model. We found 16p11.2dels clustered significantly differently from controls at both NPC and NC stages, much more remarkable at NPC stage (Fig. 3B-C).

For carriers, 21 of 16p11.2 genes showed significantly down-expression in 16p11.2del neuron cells at both NPC and NC stages (Fig. 3E and 5A, Table S5-6, Fig. S2 for qPCR verification), and down-expression levels were more in NPCs (absolute mean log2FoldChange=1.15, 0.59-1.70) than in NCs (absolute mean log2FoldChange=0.94, 0.65 to 1.13). For NPC stage, we identified 2263 differentially expressed genes (DEGs, 590 downregulated and 1673 upregulated genes, Table S7). Gene ontology analysis of DEGs enriched 967 GO terms reaching significance (q<0.05). Among top ten biological processes, seven processes were related with cell morphogenesis and each contains more than 128 DEGs (including cell morphogenesis, cellular component morphogenesis, cell morphogenesis involved in differentiation, cell part morphogenesis, plasma membrane bounded cell projection morphogenesis, cell projection morphogenesis, neuron projection morphogenesis, Fig. S3). KEGG analysis of DEGs enriched in ECM-receptor interaction, cell cycle, pathways in cancer, p53 signaling pathway, protein digestion and absorption, cytokine-cytokine receptor interaction, axon guidance and MAPK signaling pathway. were involved above pathways repeatedly (Fig. 5C). Many genes outside of the 16p11.2 locus were down-expressed, in which 69 were constrained gene (pLI>0.99), and twenty genes (*DPYSL5, KIF5C, CACNA1E, EBF3, CLCN4, GRIN2D, RELN, GRIN2A, LGI1, ASXL3, DCAF8, PDE4D, NFASC, AFF3, APC2, KIF5A, CACNA1B, RUSC2, NRG1, KCNMA1, DCAF8, NEXMIF*) were definite NDD genes with autosomal dominant inheritance model.

We performed weighted gene co-expression network analysis (WGCNA, WGCNA package v. 1.70-3) in R (v.4.1.2) (Langfelder and Horvath, 2008) to identify the pattern of gene expression correlated with NDD features. Eight modules were detected in NPCs, of which six modules had significant associations with at least one clinical variable analyzed and three modules were significantly related to NDD (Fig. 3F). NPC-MEmaroon had the most significantly negative correlation with several childhood neurodevelopmental phenotypes, including NDDs (p=5e-05), ID (p=3e-03), ASD (p=3e-03), seizure (p=2e-02), learning difficulty (LND, p=3e-03) and language delay (LGD, p=3e-03) (Genes were listed in the Table S9). Significantly, several genes of 16p11.2 locus including *MAPK3, TAOK2* and *PRRT2* were the top nodes in NPC-Memarron module (Fig. 3G). The correlation between gene significance and module membership was moderate (cor=0.56, p=1.1e-130). We further performed biological function for genes in NPC-Memarron module. As a result, the first ten extracted GO terms are associated with neuron development (Fig. 3H), included neuron projection development, synaptic signaling, trans-synaptic signaling, chemical synaptic transmission, anterograde trans-synaptic signaling, cell projection morphogenesis, cell part morphogenesis, plasma membrane bounded cell projection morphogenesis, neuron projection morphogenesis and cell morphogenesis involved in neuron differentiation. Five 16p11.2 genes were included, in which *MAPK3 and TAOK2* involved in six terms, and *PRRT2, KCTD13*, and *DOC2A* were in four terms (Fig. 3H). Besides, the top enriched terms contain some genes outside of 16p11.2 and critical for neurodevelopment, such as *ACHE, STXBP1, YWHAG, VAMP2, CACNA1E, DLG4, KCNQ3, SYT1* and *GNAO1*. KEGG analysis of NPC-Memarron module also showed several pathways related to neuron development including axon guidance and neurotrophin signaling pathway (Fig. 3I).

For NC stage, 204 DEGs were identified (98 downregulated and 106 upregulated genes, Table S8), which was much less than that of NPC stage. Pathway analysis of DEGs revealed *MAPK3* was the hub node of enriched network (Fig. 5D). WGCNA analysis detected two modules (Melightpink4, Melavenderblush2) were significantly negatively related to NDD phenotypes. This module contains 41 down-expressed genes. Genes within NC-MElightpink4 module was further analyzed as the model was most associated with NDDs (p=2e-18, Fig. 3J). The correlation between gene significance and module membership was high (cor=0.91, p=2.6e-27, Fig. 3K, gene list in Table S10) and several 16p11.2 genes were involved in the NC-MElightpink4 model (Fig. 3K). No significant pathway was enriched from DEG or GWCNA analysis. In summary, transcriptomic profiles from hiPS-derived NPCs demonstrated that 16p11.2del disrupts multiple pathways important for early neurogenesis. In addition, high down-expression of 16p11.2 genes, more DEGs (2263 *vs*. 204) and NDD-related modules (6 *vs*. 2) and pathway enriched from WGCNA were detected in NPCs than in NCs, implied these neural dysfunctions are much apparent at the NPC stage.

### Larger soma and hyperactivity of mature cortical neuron cells with 16p11.2del

The differentiation capacity of iPSC into NPCs was not different between 16p11.2del carriers and controls as demonstrated by a comparable Nestin+ cell population (90.12 *vs.* 90.92%, p>0.05, Fig. 4A-B). 16p11.2dels showed significantly bigger cell body than that of controls (142.69 vs. 133.46, p=0.044, Fig. 4C-D) at the NC stage, but not the NPC stage (*vs.*181.96, p>0.05, Fig. 4C-D), which is opposed to transcriptomic profiles showing more DEGs at NPC stage than NC stage. As Multi-electrode array (MEA) recorded few spikes (0-44) at early stage of NCs, we continuously collected the weighted mean firing rates since Day9 to Day33 (Day9-33). As shown in Fig. 4E-F, the 16p11.2dels showed a significantly increased firing rates (p= 0.008) comparing with control NCs, which is consistent with what was found in previous hiPSC-derived 16p11.2del neuron cells (Deshpande et al., 2017; Sundberg et al., 2021).

**Figure 4.**
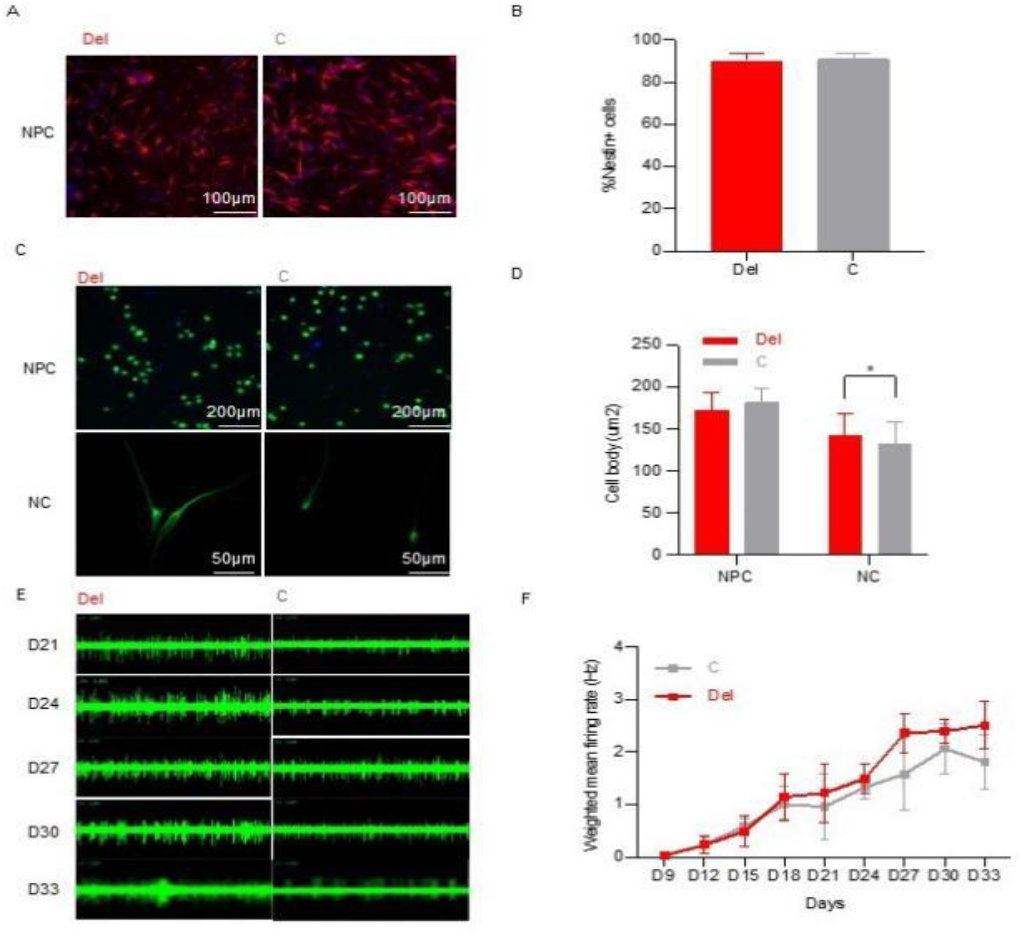
Soma size and electrophysiological properties of hiPSC-derived mature cortical neuron cells with 16p11.2del. A-B, Representative images (A) and quantification analysis (B) of differentiation efficiency of NPCs between 16p11.2del and control. The percentage of NPCs with Nestin+ were used quantify the differentiation efficiency. C-D, Representative images (C) and quantification analysis (D) of cell body size between 16p11.2del and control at NPC stage (Day0) and NC (Day7) stage. The body size of NPCs between two groups were compared by living cell staining in CELLINSIGHT CX5 (Thermo scientific) with 96-well plate, and the inoculum quantity was 1*104/well. C: N=6; Del: N=8. The body size of early NCs between two groups were compared after MAP2 staining. C: 43 cells; Del: 118 cells. T test with two tailed p values, * p<0.05. E-F, Electrophysiological Signatures of NCs recorded by MEA from Day9 to Day33 after plating NPCs. Representative images of electrophysiological properties (E) and quantification analysis (F) of weighted mean firing rates between 16p11.2del and control. X axis shows timeline after plating. The cumulative differences of weighted mean firing rates of NCs from Day9 to Day33 was compared. C: N=3; Del: N=5.

**Figure 5.**
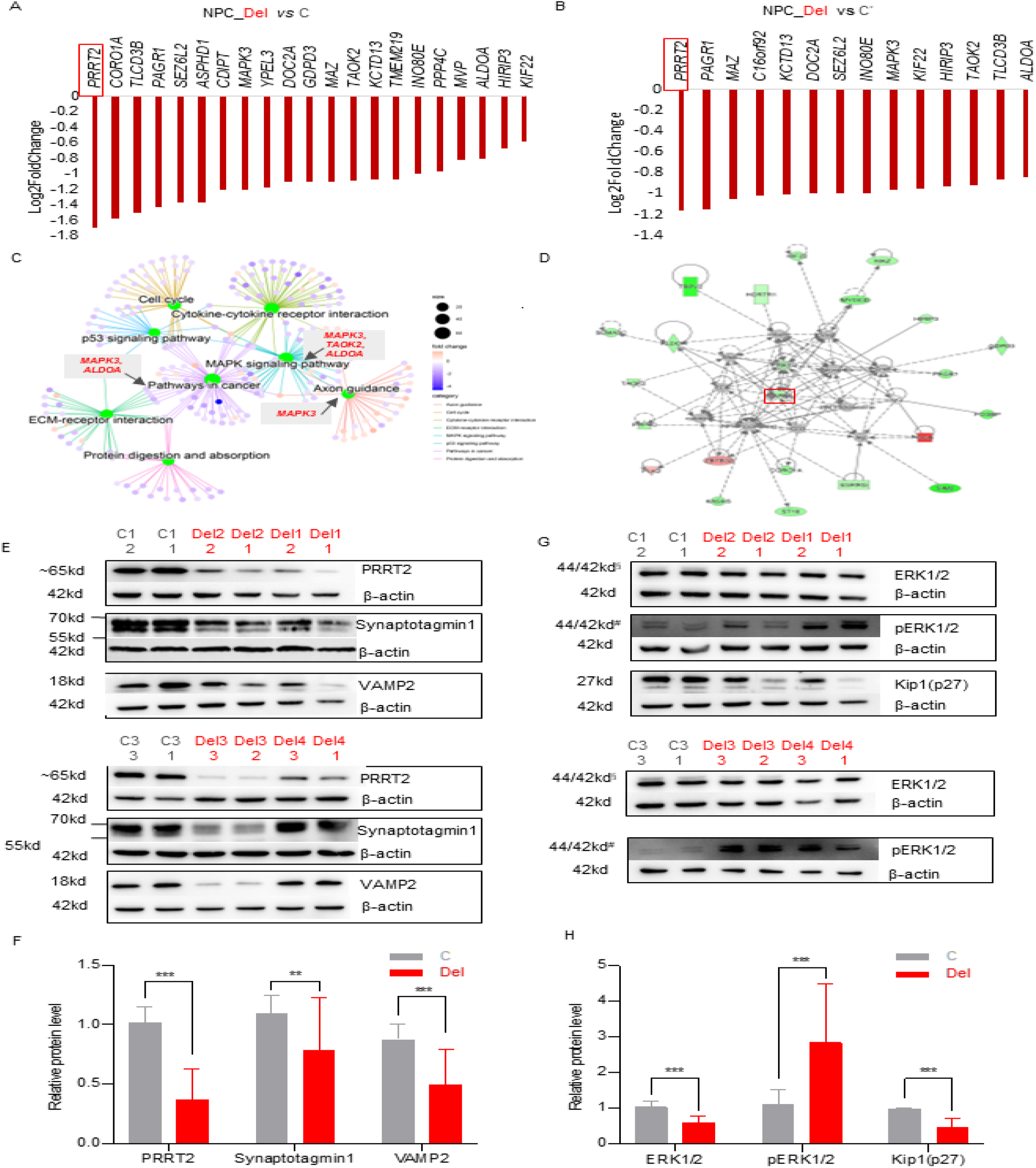
*MAPK3* emerges as driver signal of 16p11.2del at both transcriptional and protein levels resulting to over-active ERK signal in cortical neuron cells. A-B, 16p11.2 genes with down-expressions were ranked by Log2FoldChange. PRRT2 (red box) was the top gene with largest Log2FoldChange in our NPCs (Del: N=10; C: N=10, A) and in previous hiPSC-derived 16p11.2del NPCs(Roth et al., 2020) (Del: N=13; C: N=7, B). C-D, Enrichment analysis of differentially expressed genes (DEGs) between 16p11.2del and controls in NPC (C) and NC stages (D). The gene in the central network of DEGs, MAPK3, is marked by the red box. E-H, Western analysis of PRRT2 (E-F), MAPK3 (ERK1) (G-H) and other related proteins between 16p11.2dels (Del: N=8) and sex-matched controls (C: N=4). The number below sample represents for clone ID. The experiment was repeated three independent times for each clone. All values were normalized by β-actin and reported as Foldchange. T test with two tailed p values, *** p<0.001, ** p<0.01, * p<0.05.

The asynchronous results between transcriptomics, cellular morphology and electrophysiological properties of 16p11.2del neuron cell at two developmental stages suggested the molecular dysfunctions occurred prior to cellular phenotypes in neural progenitor cells, acting as primer events for altered morphological or electrophysiological properties in mature neuron cells.

### *MAPK3* emerges as driver signal related to abnormal developmental in in cortical neuron cells with 16p11.2del

We tried to explore the driver signal/gene of 16p11.2del contributing to multiple pathway dysfunction during neuron development unbiasedly using the prioritization of 16p11.2 gene based on their roles in enriched functional pathway and the foldchange levels. Frist, enrichment analysis of DEGs between 16p11.2del and control NPCs reveals *MAPK3* as the central hub of disrupted transcriptomic profiles (Fig. 5C). Second, the biological function for genes in NPC-Memarron module revealed *MAPK3* signal enriched in multiple pathways related to neuron/axon developmental function pathways (Fig. 3H and 3I, gene list in Table S11 and Table S12). *MAPK3* ranked as top seventh down-expression gene in NPCs, and showed significantly down-expression than NCs (Fig. 5A). For 16p11.2del NCs, although we failed to detect the biological function for genes in NC-MElightpink4 module, IPA analysis still detected *MAPK3* as the hub of network dysfunction. Gene in the NC-MElightpink4 module also included MAPK3 in two pathways not reach to significance (axon guidance, MAPK signaling pathway).

We performed replication analysis using the published raw transcriptomic data of cellular models derived from hiPSC with 16p11.2del, including cortex neurons (Roth et al., 2020), dopaminergic neuron (Sundberg et al., 2021), and cortical organoids (1M and 3M refers to 1 Month and 3M organoid) (Urresti et al., 2021). For the neuron cell data (Roth et al., 2020), DAVID gene enrichment analysis showed *MAPK3* was involved in 60 out of the 73 enrichment categories. Notably, they pointed out impacted MAPK signal based on their WGCNA (see their Table S6). For dopaminergic neuron cell data (Sundberg et al., 2021), WGCNA detected two modules that were highly corrected with 16p11.2del, and the ingenuity pathway analysis of Module 19 showed ERK pathway was central in the gene network (see their Fig. S4). For the cortical organoid data (Urresti et al., 2021), although the sampling timepoint were different from that of our NCs (1 month old), significant down-expression of *MAPK3* was observed at both transcriptomics and proteomics profile. These replication analyses confirmed the driver signal of *MAPK3* to pathways dysfunction related to neuron development.

Besides *MAPK3, PRRT2* is worthwhile mentioning because *PRRT2* had the biggest foldchange value than other 16p11.2 gens (Fig. 5A, qPCR validation in Fig. S2). The GO analysis of MEmarron module showed *PRRT2* involve four of top pathways of neuron development (Fig. 3H). We also evaluated the fold change values of 16p11.2 genes were reported only in hiPSC-derived cortex neuron cells, showing *PRRT2* ranked as the top down-expression genes (Fig. 5B) (Roth et al., 2020).

We further tested the consequence of *MAPK3 and PRRT2* haploinsufficiency at NPC stage. For MAPK3 and its downstream molecules (Fig. 5G-H), western blotting showed the 16p11.2dels had significantly decreased ERK1 levels (0.6 *vs.* 1.04, p<0.001), but increased phosphorylated-ERK1 (pERK1) levels (2.84 *vs.* 1.11, p<0.001) compared to the controls, indicating aberrant up-regulation of ERK activity due to 16p11.2del. Besides, Kip1(p27) protein, the direct targets of the ERKs, showed significantly decreased expression (0.46 *vs.* 0.99, p<0.001) in the 16p11.2delss, which was consistent with previous results of 16p11.2del mouse (Pucilowska et al., 2015). The proteins related to PPRT2, including synaptotagmin1 and VAMP2, showed signifyingly decreased protein levels in 16p11.2dels than controls (PRRT2, 0.37 *vs.* 1.02, p<0.001; Synaptotagmin1, 0.79 *vs.* 1.09, p=0.006; VAMP2, 0.5 *vs.* 0.87, p<0.001, Fig. 5E-F).

### *MAPK3/PRRT2* overexpression reversed abnormal ERK signal and hyperactivity in cortical neuron cells with 16p11.2del

To provide further evidence of driver signal of *MAPK3* and *PRRT2*, we performed rescue study at Day-3 using the lentiviral vectors (Blue font in Fig. 2C), and tested the expression of related proteins on Day0, recorded the electrophysiological properties since Day9 to Day33 (Day9-33). With two lentiviral dosages of *PRRT2* ((R (+) and R (++)), the levels of *PRRT2*-related proteins, including Synaptotagmin1 ((R (+): p=0.001; R (++): p=0.01) and VAMP2 ((R (+): p=0.003; R (++): p=0.004), were significantly increased (Fig. 6A-B). Overexpression of *MAPK3* also significantly reversed the activity of p-ERK1 ((R (+): p=0.001; R (++): p=0.01) and increased the expression of Kip1(p27) ((R (+): p<0.001; R (++): p<0.001, Fig. 6C-D). High weighted mean firing rates (Day9-33) of 16p11.2del NCs were significantly reversed by the reintroduction of *PRRT2* (p=0.003, Fig. 6E-F) and *MAPK3* (p=0.0001, Fig. 6G-H). In addition, more reversed ratio of electrophysiological properties was noted for *MAPK3* lentiviral vectors than for *PRRT2* lentiviral vectors (Fig. S4), suggesting 16p11.2del NCs benefit more from *MAPK3* than *PRRT2* overexpression.

**Figure 6.**
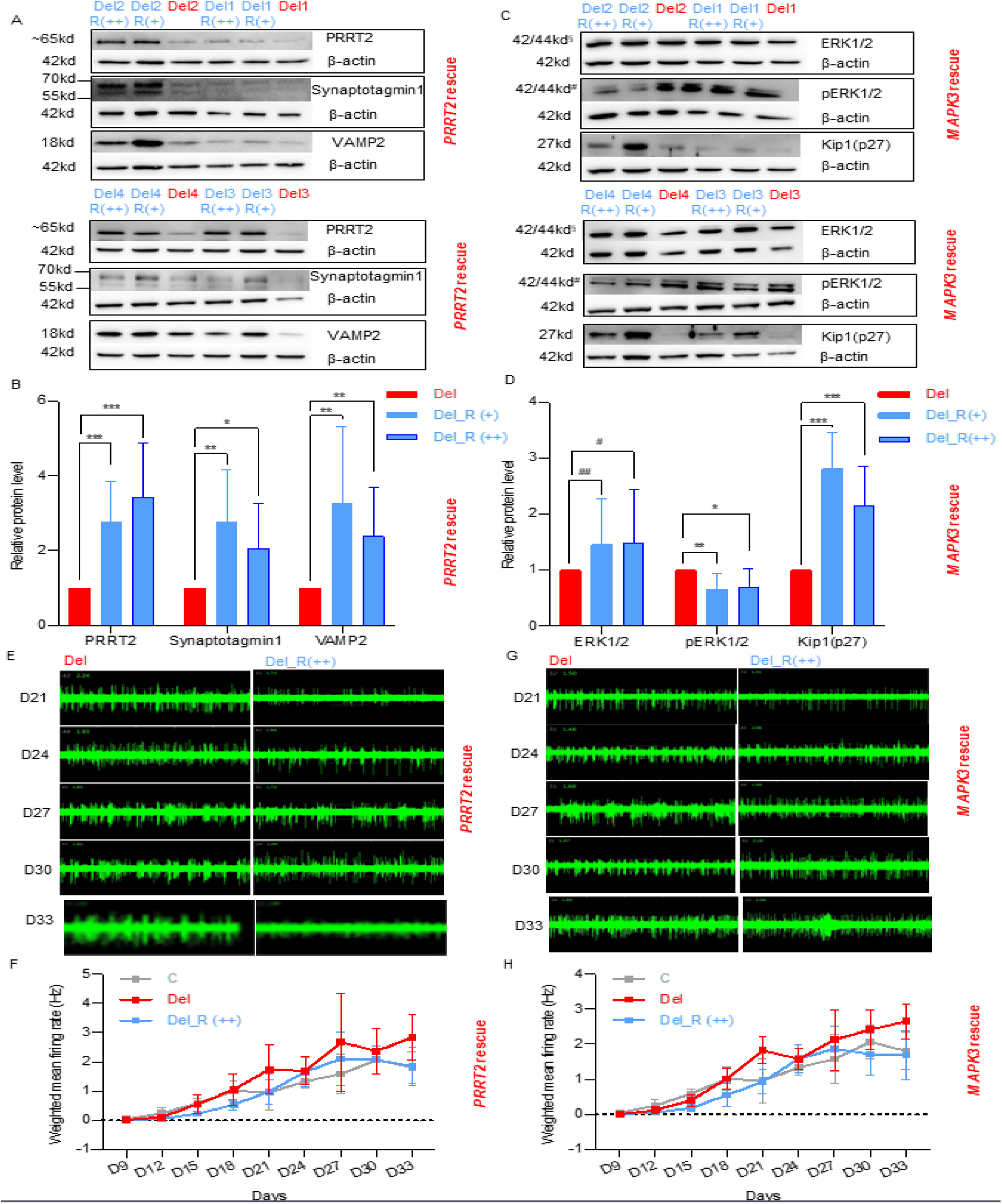
*MAPK3/PRRT2* overexpression reversed abnormal ERK signal and hyperactivity in cortical neuron cells with 16p11.2del. A-D, Increased protein expressions (NPC stage) and reduced electrophysiological properties (NC stage) in 16p11.2del neuron cells treated with PRRT2 lentiviral. A-B, Western blotting (A) and quantification analysis (B) of PRRT2, Synaptotagmin1 and VAMP2 in 16p11.2del NPCs treated with PRRT2 or blank lentiviral. C-D, Representative images of electrophysiological signals recorded by MEA (C) and quantification analysis of weighted mean firing rates (D) in 16p11.2del NCs treated with PRRT2 lentiviral and blank lentiviral. X axis shows timeline after seeding NPCs. E-H, Increased protein expressions and reduced electrophysiological properties in 16p11.2del NPCs treated with MAPK3 or blank lentiviral. E-F, Western blotting (E) and quantification analysis (F) of ERK1, pERK1 and Kip1(p27) in 16p11.2del NPCs treated with MAPK3 or blank lentiviral. G-H, Representative images of electrophysiological activity recorded by MEA (G) and quantification analysis of weighted mean firing rates (H) in 16p11.2del NCs treated with MAPK3 lentiviral and blank lentiviral. For Western blotting, four 16p11.2del clones were picked for rescue assay. Each cell clone was treated with two lentiviral dosages, and with PRRT2/MAPK3 lentiviral and blank lentiviral as well (Del). R (+) represents for low dosage (2.4×104 TU for PRRT2 and 2×104 for MAPK3) and R (++) represents for high dosage (1.2×105 TU for PRRT2 and 1×105 for MAPK3). Dosage of blank was 1.2×105 TU for PRRT2 and 1×105 for MAPK3 experiment. For electrophysiological properties, five 16p11.2del clones were picked for rescue assay. Each cell clone was treated with PRRT2 (1.2 105 TU)/MAPK3 (1×105 TU) lentiviral and blank lentiviral (Del). Experiments was repeated three independent times with similar results. All values were normalized by β-actin and reported as Foldchange. T test with two tailed p values, *** p<0.001, ** p<0.01, * p<0.05.

### Residual haplotype-specific *MAPK3* expression, transcriptomics profile and electrophysiological properties in cortical neuron cells with 16p11.2del

We noticed some 16p11.2 genes showed different expressions between intra-family carriers. In view of remaining haplotype, six genes of 16p11.2 interval (*PRRT2, PAGR1, GDPD3, SEZ6L2, YPEL3* and *MAPK3*, see Table S13 and Fig. S4) showed significantly lower expression in Del_H1 NPCs (n=5) than Del_H2 NPCs (n=5). However, the qRT-PCR and WB only validated significant down-expression of *MAPK3* in Del_H1, at both RNA (0.26 *vs.* 0.58, p=0.039, Fig. 7A) and protein level (0.48 *vs.* 0.73, p<0.001, Fig. 7B). The transcriptomic profiles between two haplotypic subgroups were further compared using DEseq2, identifying 377 DEGs at NPC stage. GO analysis showed that DEGs is still enriched for terms related to synaptic development and function, including synaptic signaling, chemical synaptic transmission, anterograde trans-synaptic signaling, signal release, modulation of chemical synaptic transmission, regulation of trans-synaptic signaling, neurotransmitter transport, vesicle-mediated transport in synapse, neurotransmitter secretion (Fig. 7C).

**Figure 7.**
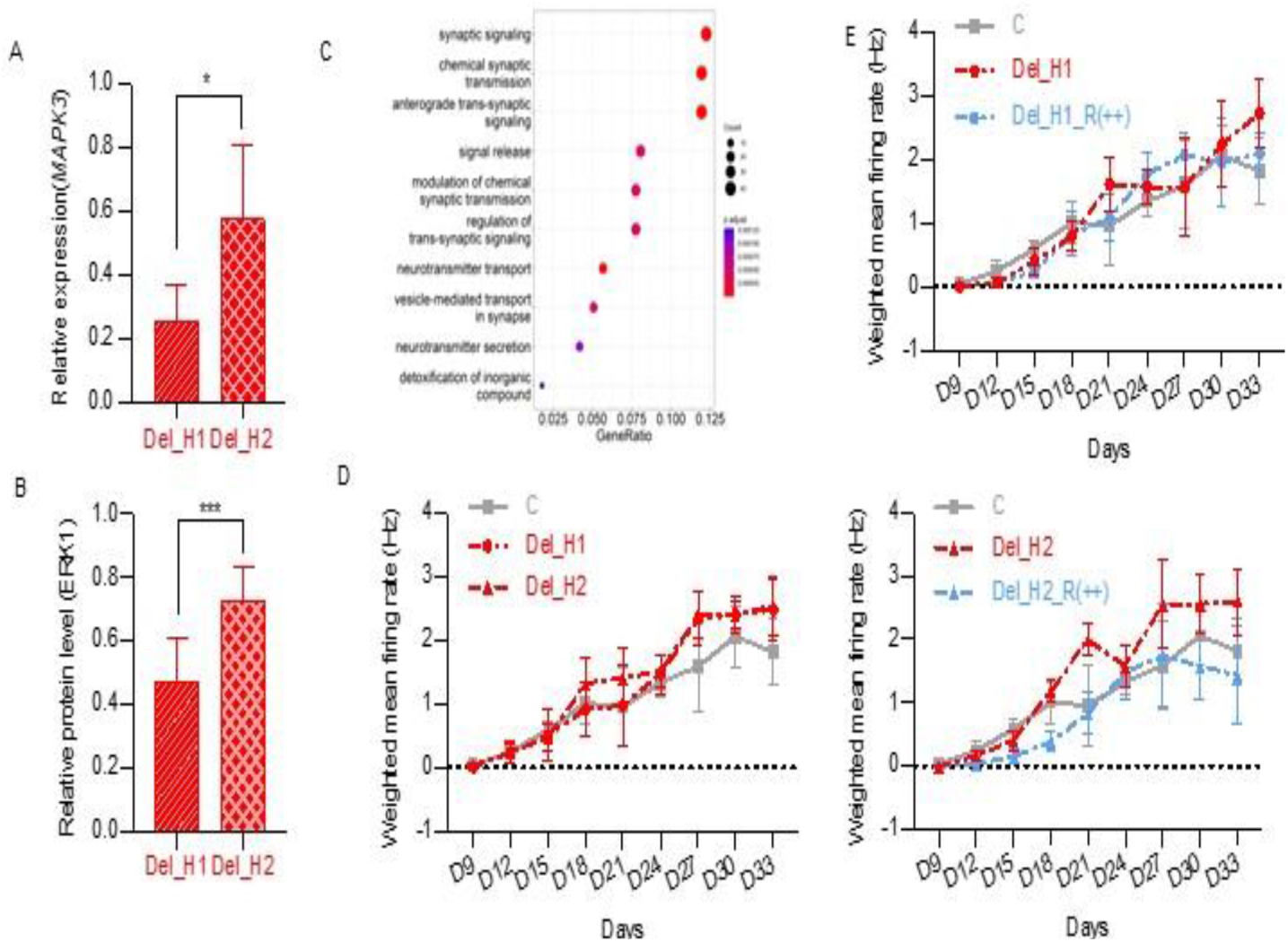
Residual haplotype-specific *MAPK3* expression, transcriptomics profile and electrophysiological properties in cortical neuron cells with 16p11.2del. **A,** Quantification of *MAPK3* mRNA levels in 16p11.2del NPCs carrying different residual haplotypes with qRT-PCR experiment. Del_H1: N=4, Del_H2: N=5. **B,** Quantification of ERK1 protein in 16p11.2del NPCs carrying different residual haplotypes. Del_H1: N=4. Del_H2: N=4.**C,** GO biological processes enriched with DEGs for 16p11.2del NPCs carrying different residual haplotypes. Del_H1: N=5. Del_H2: N=5. **D**, Quantification of weighted mean firing rates in 16p11.2del NCs carrying different residual haplotypes. Del_H1: N=2, Del_H2: N=3. X axis shows timeline after seeding NPCs. **E**, Quantification of weighted mean firing rates in different 16p11.2del NCs treated with *MAPK3* (1×10^5^ TU). X axis shows days after seeding NPCs. Del_H1 with *MAPK3* lentiviral: N=2. Del_H1 with blank lentiviral: N=2. Del_H2 with *MAPK3* lentiviral: N=3. Del_H2 with blank lentiviral: N=3. T test with two-sided p value less than 0.05 was considered as statistically significant tailed p values, * p<0.05. *** p<0.001.

**Figure 8.**
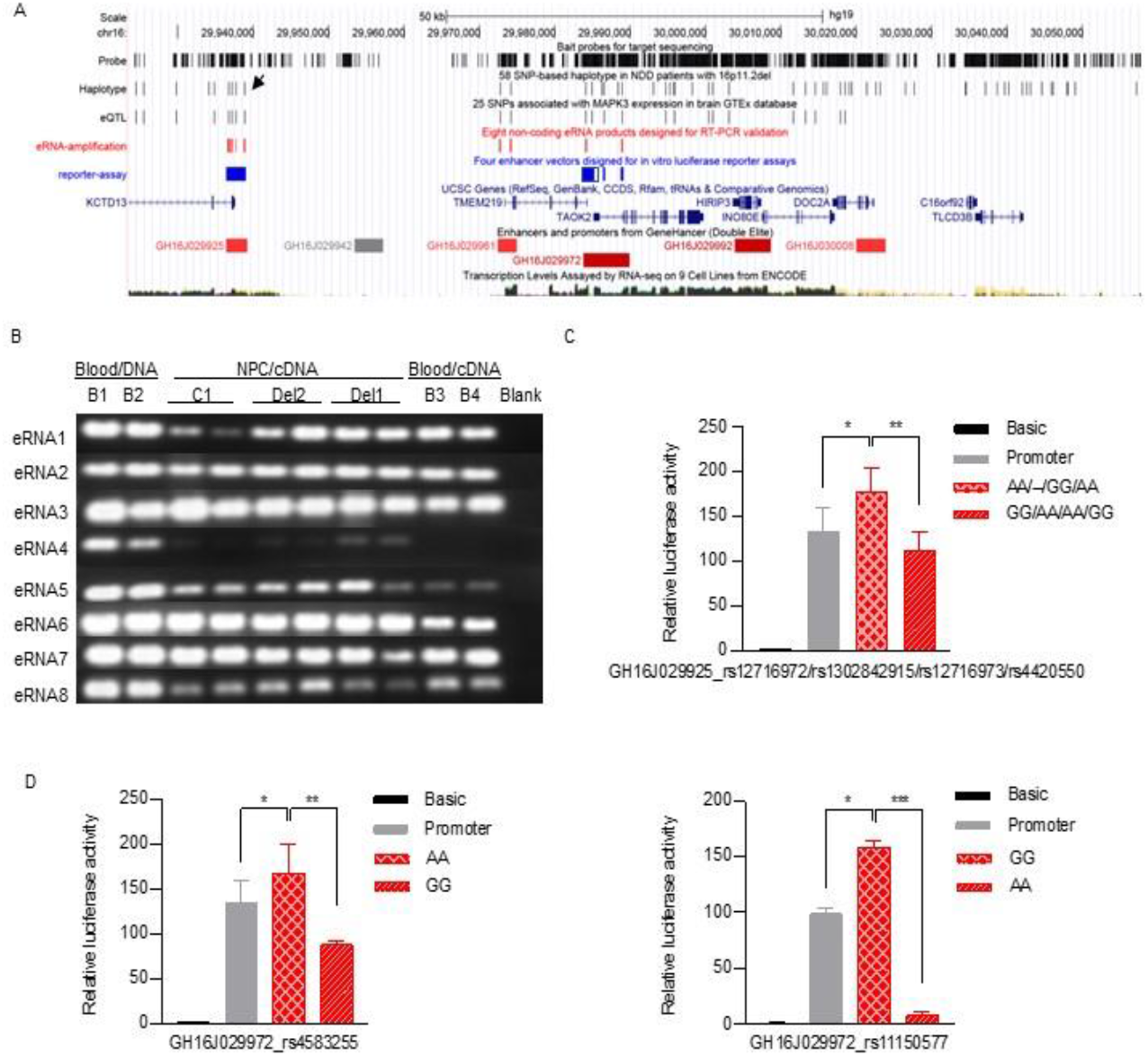
The cis-regulation of *MAPK3* haploinsufficiency in 16p11.2 deletion due to non-coding variant. **A,** 16p11.2 region (chr16:29,923,426-30,057,770, hg19 version) showed gene content, capture probes for target sequencing, 58 SNPs-based residual haplotype (black tracks), 25 SNPs associated with *MAPK3* expression in brain GTEx database (black tracks), eight non-coding eRNA products designed for RT-PCR experiments (red tracks), four enhancer fragments designed for *in intro* luciferase reporter assays (blue tracks). One reported regulatory SNP associated with schizophrenia was labeled as black head-arrows. One failed amplicon of enhancer fragment was labeled as asterisk. **B,** RT-PCR products of eight non-coding eRNAs using cDNA template of NPCs (N=6, C1, Del1 and Del2) and cDNA template of Blood ((N=2, B2 and B4). The PCR product of genomic DNA template of blood (N=2, B1 and B2) was used as controls to confirm unique genomic sequence. The primers of eight non-coding eRNA products were listed in Table S17. **C,** Comparison of the luciferase activities of entire GH16J029925 (containing four haplotype-specific enSNPs, rs12716972/rs1302842915/rs12716973/rs4420550. AA/--/GG/AA as major enhancer and GG/AA/AA/GG as minor enhancer) in 293HEK cell-lines). **D,** Comparison of the luciferase activities of caudal part of GH16J029925 (two haplotype-specific enSNPs) in 293HEK cell-lines. Left panel: rs4583255, AA as major SNP and GG as minor SNP; Right panel: rs11150577, GG as major SNP and AA as minor SNP. The Y-axis values represent fold changes of luciferase activity relative to the empty pGL3 promoter vector. T test with two tailed p values, *** p<0.001, ** p<0.01, * p<0.05.

We further performed replication analysis using the published transcriptomic data of hiPSC-derived 16p11.2del neuron cells. Among ten such cells Julien G Roth et al studied (Roth et al., 2020), only one (Sample ID: DEL_10; SFARI Subject ID: 14799.x1) was phased as Del_H1, while other as Del_H2 based on their genotypes (Simons Vip, 2012). The transcriptomic data of three clones of Del_H1 (Sample DEL_10) was compared with three clones of one sex-matched Del_H2 (Sample ID: DEL_9, SFARI Subject ID: 14824.x13) (see Table S14 for their relative expression of 16p11.2 interval genes). Consistent with our observation, the expression of *MAPK3* (p=0.0000696, Fig. S5) along with *KIF22* (p= 0.0000163) were significantly lower in Del_H1 than that in Del_H2.

We also tried to compare the cellular morphology and electrophysiological properties of different 16p11.2del NCs. After differentiated to Day23, only five 16p11.2del NC clones (Del_H1: n=2; Del_H2: n=3) maintained normal development and were available for the comparison. We found the weighted mean firing rates (p=0.11 for Day9-33, Fig. 7D), body size (p=0.16, Fig. S6) were similar between two groups. More important, *MAPK3* overexpression completely reversed the hyperactivation of NCs genotyping as Del_H2 (p=0.0001 for Day9-33, Fig. 7E), but failed to significantly reverse the hyperactivation of NCs genotyping as Del_H1(p=0.42, Fig. 7E). The comparison of transcriptomics profile, electrophysiological properties and rescue experiment in different 16p11.2del neuron cells suggested the minor residual haplotype (Del_H1) is associated with severe *MAPK3* down-expression and worsen cellular phenotype in.

### Enhancer SNPs contribute to residual haplotype-specific *MAPK3* expression via cis-regulation

Enhancers are regulatory DNA sequences and able to modulate the expression of target gene dozens of kb away (Levo and Segal, 2014). Expression quantitative trait locus (eQTL) analysis is a powerful method to identify a link between the expression of a target gene and the enhancer sequences (Gibson et al., 2015). In order to explore the genetic root of *MAPK3* expression variation detected inhiPSC-derived 16p11.2del NPCs, we extracted 135 SNPs which are significantly associated with *MAPK3* from Genotype-Tissue Expression project (GTEx, https://www.gtexportal.org/home) (Table S15), and intersected with 58 SNPs of the residual haplotype. Among them 25 SNPs were significantly associated with *MAPK3* expression in the brain eQTL (Fig. 1D and Fig. 8A, labeled yellow in Table S1), with down-expression for all minor alleles (Table S16, Fig. S7). This regulation is consistent with our observations in the 16p11.2del NPCs, where *MAPK3* down-expression is associated with minor residual haplotype.

None of these 25 eQTL SNPs is coding variant. As the residual haplotype (chr16:29936654-29988941) and *MAPK3* (chr16:30125426-30134630) is 137kb far away and locate in different LD blocks (Fig.1C), we suspected that SNPs on residual haplotype exert cis-regulatory effects via long-range spatial contacts. We further analyzed the locations of regulatory elements in 16p11.2 region (GeneCard database) (Fishilevich et al., 2017) against the 25 SNPs, and detected ten SNPs mapped to three regulatory elements (GH16J029925, GH16J029961, GH16J029972, Fig.8A), with 2-4 SNPs in each element (see Table S1 for detailed location of ten regulatory SNPs), suggesting ten SNPs may work as enhancer SNPs (enSNP) to regulate *MAPK3* expression.

The identification of enhancer RNA (eRNA) product could be the direct evidence of enhancer (Andersson et al., 2014). First, we aligned RNA-Seq data of NPCs for the genomic locations of enhancers. RNA read piled up at non-coding regions of three putative enhancers (Fig. S8), confirming the hypothesized eRNA. The eRNA levels in the 16p11.2dels were about half of that of controls. No significant difference was seen in the 16p11.2dels with different (major or minor) residual haplotypes. We also performed RT-PCR experiment after DNA digestion to validate the existences of eRNA fragments (130-271bp, primer name and location in Table S17) (red vertical bar-lines in Fig. 8A). Eight small eRNA fragments covering intronic region (Lane 1-8 in Fig. 8B) were successfully amplified and confirmed by Sanger sequencing. Six eRNAs (eRNA1-3, 5, 7,8) expressed in both NPCs and blood, while two eRNAs (eRNA4 and 6) express only in NPCs. Reads covering these success amplicons were also seen in RNA-Seq (Fig. S8).

To illuminate the regulatory role of the haplotype composed of multiple enSNPs, we performed *in vitro* luciferase assays for two enhancers (GH16J029925 and GH16J029972, covering eight SNPs) after amplifying haplotype-specific and long enhancer fragments (primer name and location were listed in Table S17). One enhancer fragment (1627bp) covering cranial part of GH16J029972 (black box in Fig. 8A, labeled yellow in Table S17, rs11901/rs4318227) was failed in PCR amplification. Remained three enhancer fragments were constructed into pGL3-promoter vector (blue vertical bar-lines in Fig. 8A), including one long fragment (2564bp) covering four enSNPs (rs12716972/rs1302842915/rs12716973/rs4420550) and entire GH16J029925, two short fragments (326bp and 367bp) covering two enSNPs (rs11150577 and rs4583255) and caudal part of GH16J029972.

The enhancer activities of different enhancer fragments on gene expression were compared in HEK293T cells. As shown, minor GH16J029925 [GG/AA/AA/GG of rs12716972/rs1302842915/rs12716973/rs4420550] led to significantly reduced luciferase activities than major GH16J029925 [AA/--/GG/AA] (Fig. 8C, p=0.002). Meanwhile for caudal part of GH16J029972, minor enSNPs [AA of rs11150577, GG of rs4583255] led to signifyingly reduced luciferase activities than major enSNPs [GG of rs11150577, AA of rs4583255] (Fig. 8D, rs11150577, p=0.001; rs4583255 p=0.005). The reduction index of rs11150577 was higher than that of rs11150577 (94.6% *vs.* 47.4%). These results proved that residual haplotype-specific SNPs on enhancer (GH16J029972 and GH16J029925) can regulate the expression of *MAPK3* by enhancer RNA product

### Minor residual haplotype of 16p11.2 locus is associated with NDD/ASD rather CS phenotype in 16p11.2del carriers

We quested whether the residual haplotype of 16p11.2 locus is associated with NDD clinical phenotypes. As the quantitative severity of NDD phenotype was not available for some of the 16p11.2del carriers, we compromised to compared the frequency of minor residual haplotypes in three different cohorts (Table 1), including 18 ASD patients with 16p11.2del from Simons Foundation Autism Research Initiative (SFARI Base, https://www.sfari.org/, SFARI-ASD-16p11.2del), 29 CS patients with 16p11.2del from Peking Union Medical College Hospital (PUMC-CS-16p11.2del) and our NDD cohort. We speculated that PUMC-CS-16p11.2del patients are prone to be non-NDD or mild NDD because they came to hospital for CS rather NDD. The residual haplotype of ASD and CS patients were phased by whole genome sequencing or SNP array. The minor residual haplotype was significantly enriched in NDD cohort than in CS cohort (61% *vs.* 10.31%, p<0.001), but similar between NDD and ASD cohort (61% *vs.* 44.4%, p>0.05). The frequency of minor residual haplotype in the NDD and ASD cohorts were also deviated from that of general population (34-39%). The finding from this small sample size suggested possible association between the residual haplotype and variable NDD phenotypeof 16p11.2del.

## DISCUSSION

Recurrent 16p11.2del is associated with broad NDDs with differential penetrance (Weiss et al., 2008; Steinberg et al., 2014; Hudac et al., 2020; Rein and Yan, 2020). 16p11.2del are also associated with other disorders beyond neurological system, such as congenital scoliosis (Wu et al., 2015), BMI (Jacquemont et al., 2011), neuroblastoma (Egolf et al., 2019), and anemia/iron deficiency via modifier copy of *BOLA2* (Giannuzzi et al., 2019). Recent, two groups revealed 16p11.2del is risk of congenital anomalies of the kidney and urinary tract (CAKUT) (Verbitsky et al., 2019; Yang et al., 2020). Both studies pointed out that *TBX6* gene dosage is a major determinant of the pathogenesis of CAKUT when it combines with three known hypomorphic alleles of CS (rs2289292T-rs3809624C-rs3809627A) our group studied before (Wu et al., 2015).

The emergence of hiPSC technology offers an informative tool to explore the underlying mechanisms of NDD-related CNVs, In this model, both genomics information, transcriptomics profile and cellular morphology and electrophysiological properties of cortical neurons are available, allowing us analyzed the pathophysiology of interested variant at the cellular level (Drakulic et al., 2020), and hiPSC-derived neuron cell models have been created for 7q11.23 (Adamo et al., 2015), Kleefstra syndrome (Frega et al., 2019) and 22q13 deletion syndrome (Shcheglovitov et al., 2013). hiPS-derived neuron model of 16p11.2del have been generated (Deshpande et al., 2017; Roth et al., 2020; Sundberg et al., 2021; Urresti et al., 2021). However, these models were created from Western 16p11.2del cohort and sorted in Simon Foundation, no resource was available for Asian population. We generated 20 hiPSC lines for two Chinese familiar 16p11.2del, and banked them as publicly available resource (China Center for Type Culture Collection, CCTCC, http://www.cctcc.org) for exploring the ethnic difference of 16p11.2del in future.

In general, genes within CNV are recognized as potentially causation for specific clinical phenotypes, with haploinsufficiency in deletion and triplosensitivity in duplication (Harel and Lupski, 2018), for example, *ELN* of 7q11.23 deletion (Williams-Beuren syndrome) or *TBX1* of 22q11.21 deletion. Previous studies of the peripheral blood (Blumenthal et al., 2014), patient-derived or CRISP-edited isogenic iPSCs (Tai et al., 2016) and iPSC-derived neuron of 16p11.2del (Roth et al., 2020) had confirmed the pathogenicity of 16p11.2. However, the drive gene of NDDs in 16p11.2del is unknown so far although three candidate genes including *MAPK3, KCTD13, TAOK2* have been considered from Drosophila, mouse and zebrafish models(Golzio et al., 2012; Portmann et al., 2014; Arbogast et al., 2016; Iyer et al., 2018; Pucilowska et al., 2018; Richter et al., 2019), but is failed to be confirmed consistently in human iPSC model (Zufferey et al., 2012; Pucilowska et al., 2015; Escamilla et al., 2017; Arbogast et al., 2019).

Herein, we successfully differentiated ten hiPSCs of four 16p11.2del carriers to the cortical neuron cells. Using the transcriptomic profile and cellular phenotypes of hiPSC-derived neurons at different differentiational stages, we revealed 16p11.2del resulted multiple pathway dysfunction related to neuron development, larger body and hyperactivity of neuron cells as previous studies of hiPS-derived NPC of 16p11.2del reported (Deshpande et al., 2017; Roth et al., 2020; Sundberg et al., 2021). In addition, transcriptomic abnormality occurs early in neural progenitor cells, acting as primer events for cellular morphologies and electrophysiological properties of mature neuron cells, which has not been pinpointed before. Further, the driver gene of 16p11.2 were studied unbiasedly by the integrated analyses of the foldchange of 16p11.2 gene, pathway enrichment of transcriptomic profile and rescue experiment. We concluded that *MAPK3* presented as driver signal of 16p11.2del contributing to abnormal transcriptomic profile and cellular phenotypes, from both RNA and protein levels, at both NPC and NC stages. This drive signal of NDD phenotype in 16p11.2del is much different from that of non-NDD phenotype, for example CAKUT/CS (*TBX6*) (Wu et al., 2015; Verbitsky et al., 2019; Yang et al., 2020) or anemia/iron deficiency (*BOLA2*) (Giannuzzi et al., 2019).

*MAPK3* (mitogen-activated protein kinase 3) is key component of Ras/MAPK pathway, and it is associated with ERK (extra-cellular signal-regulated kinase) pathway activation affecting neural differentiation and proliferation (Thomas and Huganir, 2004). Both the functional studies from mice (Pucilowska et al., 2015) (Blizinsky et al., 2016) and *Drosophila* models (Park et al., 2017) had revealed *MAPK3* is potential drive pathway associated with NDD phenotype of 16p11.2del. Abnormal *MAPK3* signal has been enriched from previous transcriptomic data of hiPS-derived neuron cells and organoid from 16p11.2del carriers presenting macrocephaly (Deshpande et al., 2017; Roth et al., 2020; Sundberg et al., 2021; Urresti et al., 2021).In our study, four 16p11.2del carriers presented normal head size. Enriched *MAPK3* signal from patients with different head size stressed the universal role of MAPK3 signal to 16p11.2del. Recent, *MAPK3* has been identified as risk gene of NDDs from one larger-size whole exome sequencing study of Western NDD cohort. Author stated that unstable MAPK3 protein due to mutation result into *MAPK3* haploinsufficiency (Coe et al., 2019). This association study supplied direct genetic evidence between *MAPK3* haploinsufficiency and NDD.

Yet, considering the absolute decreased foldchange values of 16p11.2 genes at NPC stages, *MAPK3* ranked as the eighth gene of 16p11.2 region (Fig. 5A, see Fig. S2 for qPCR verification), implying the contribution of other 16p11.2 genes to *MAPK3* signal. Other 16p11.2 genes have been reported to connect to *MAPK3* signal, for example, *MAPK3* and *TAOK2* were involved in synaptic assembly and signaling (Betancur et al., 2009), *ALDOA* is members of MAPK3 signal during neuron differentiation (von Kriegsheim et al., 2009). The pervasive genetic interaction between 16p11.2 genes and *MAPK3* has been reported in *Drosophila* model (Iyer et al., 2018). They detected both suppressor and enhancer, additive interaction of *MAPK3* and other genes, showing more severe phenotype of fly with double knockdown of *MAPK3^rl^* combining with either *PPP4C ^pp4-19C^* or *TAOK2 ^dTao-1^, ASPHD1 ^asph^, KIF22 ^klp68D^*. Our pathway enrichment analyses also detected *TAOK2* co-occurred with *MAPK3* in multiple neuron developmental pathways implying its contribution to *MAPK3* signal (Fig. 3H, 3I, 5C), and *TAOK2* is important molecular of *MAPK3* signal with additive role (Fig. 3I). We also found *PPP4C, KIF22* interact with MAPK3 (Fig. 5D), However, the suppressor or enhancer role of *PPP4C, KIF22* to MAPK3 signal during human neurogenesis was not analyzed, which need systemic rescue study of hiPSC-derived 16p11.2del NPCs in the future.

*PRRT2* encodes the proline-rich transmembrane protein 2 that interact with components of the SNARE complex and the fast Ca2+ sensors synaptotagmin1/2, both of which were critical for neurotransmitter release (Valente et al., 2016). Loss-of-function mutation of *PRRT2* causes benign infantile epilepsy (BIE), paroxysmal kinesigenic dyskinesia (PKD), and paroxysmal kinesigenic dyskinesia with infantile convulsions (PKD/IC). So far, *PRRT2* haploinsufficiency in 16p11.2 syndrome, and the role of *PRRT2* during early and mature neurogenesis is unclear. However, some abnormal phenotypes in neuron with *PRRT2* knockout were similar to that of 16p11.2del, for example, loss-of-function mutation of *PRRT2* resulted to increased neuronal hyperexcitability (Lee et al., 2012), *Prrt2* knockdown *in vivo* resulted in delayed neuronal migration and decreased synaptic density (Liu et al., 2016). We presented the important role of *PRRT2* haploinsufficiency during NPC stage based on its top Log2FoldChange from our and other iPSC-derived cortical neural model (Roth et al., 2020), and its role in MEmaroon module (Fig. 3G) and four enriched pathways of neuron development (Fig. 3H and Table S12). The contribution of *PRRT2* to 16p11.2del in mature neuron cells weaken significantly (Fig. 3K and Table S6). So, the drive contribution of *PRRT2* to 16p11.2del during early neurogenesis need further explore with more 16p11.2del carriers.

Variable expression of NDDs is unsolved issue of 16p11.2del, even for inter-family carriers, making it worthwhile for further exploration the modifier allele of NDD penetration in 16p11.2del carriers. Our previous 16p11.2-related CS study have proved the hypomorphic regulation allele of *TBX6* expression is from three SNPs rather single SNP (Wu et al., 2015). From the viewpoint from our above results, exploring the multiple rather one modifier/hypomorphic variant on NDD phenotypic heterogeneity among 16p11.2del is a suitable method. we explored the residual haplotype-specific transcriptomic profile, cell morphology and electrophysiological properties of 16p11.2del. Our results demonstrated minor residual haplotype resulted worse *MAPK3* down-expression (at both RNA and protein levels) and transcriptomic dysfunction of cortical neuron cells in 16p11.2del. Considering insignificant difference of cellular phenotype, we presumed that small 16p11.2del samples resulted to negative finding.

Long-distance cis-regulations, even cell-type-specific cis-regulation of non-coding variant have been reported in neural cells (Song et al., 2019) and human frontal lobe (Girdhar et al., 2018). We screened out ten haplotype-linked enSNPs who is associated with *MAPK3* expression in brain tissue and locate in three elite enhancers (GH16J029972, GH16J029961 and GH16J029925). Furthermore, the regulatory roles of six enSNP fragments (rs12716972/rs1302842915/rs12716973/rs4420550/rs11150577/rs4583255) of two enhancers (GH16J029972 and GH16J029925) were validated by *in vitro* assay. As the distance between these regulatory elements (chr16:29936654-29988941) and *MAPK3* (chr16:30125426-30134630) is quite far (about 137kb) and they locate in different LD block (Fig.1C), we speculate these two enhancers exerts cis-regulatory effects via long-range spatial contacts.

One SNP (rs4420550) on GH16J029925 (black arrow in Fig. 8A) has been reported to regulate *MAPK3* expression and increased risk of schizophrenia (Chang et al., 2021). The *in vitro* functional experiments (589bp) showed increased luciferase activities in 293HEK cells with major allele [A/A] compared with that with minor [G/G] allele, same to our results. Different from their design, we inserted entire GH16J029925 (2564bp, containing rs12716972/rs1302842915 /rs12716973/rs4420550) into pGL3-promotor vector for *in vitro* assay, showing higher luciferase activities in 293HEK cells with major enhancer [AA/--/GG/AA] compared with that with minor enhancer [GG/AA/AA/GG]. In addition, we confirmed the potential regulatory role of another enhancer, GH16J029972 using two short enSNP fragments (326bp for rs11150577, 367bp for rs4583255), which has not been reported before. We also consider that residual haplotype-specific cis-regulation in hiPSC-neuron of 16p11.2del maybe not limited in *MAPK3* as we presented in this study. Several 16p11.2 genes showed down-expression levels in NPCs with minor residual haplotype, including *PRRT2, PAGR1, GDPD3, SEZ6L2* and *YPEL3* although non-significance. Similarly, the schizophrenia study also demonstrated rs4420550 was associated with the expression of several 16p11.2 genes (*INO80E, KCTD13, TAOK2, ALDOA, HIRIP3*) beyond *MAPK3*, suggesting potential regulatory role of this long residual haplotype to other 16p11.2 genes. Consistently, we found the hyperactivation of NCs genotyped as of Del_H1 can’t be rescued *MAPK3* overexpression as that in Del_H2, indicating that the minor allele haplotype could regulates more than *MAPK3* (Fig.7E).

Based on our results from transcriptomic profile, cellular phenotype of cortical neuron, clinical referred phenotypes of affected patients, we bring up preliminary hypothesis schedule of phenotypic heterogeneity among 16p11.2del carriers: the haplotype of remained 16p11.2 locus can cis-regulate the expression of *MAPK3 via* regulatory SNPs, contribute to residual haplotype-specific transcriptomics profile and electrophysiological properties in cortical neuron cells with 16p11.2del, and finally to differential NDD phenotype (Fig. S9). We further tested this hypothesis using three different clinical patient cohorts with 16p11.2del (Table 1), showing minor residual haplotype added risk of NDD/ASD rather CS phenotype in 16p11.2del carries.

We also tried to apply this hypothesis schedule to explain severe ASD phenotype in one iPSC donor. As we described, donor Del1 (II:1 of Family 1) was diagnosed classic ASD, severe language deficiency, and her symptom was severe than other three Del carriers. We extracted low-frequency non-coding variants (MAF<10% of 600 Chinese WGS and Asian of gnomAD) (Table S18) of four 16p11.2de donors. Beside minor residual haplotype, Del1 donor harbors other 31 variants with low MAF (<10%) and 10 variants with MAF<5% respectively, which is much higher than other donors (2-10 rare variants, Table S18 and Fig. S10). Among, five Del1-specific SNPs are in one enhancer (GH16J029700, chr16:29711421-29715069, rs62060164/rs17640225/rs62060166/rs140785784/ rs138131145, yellow vertical line in Fig. S10), yielding another evidence that enSNPs are risk of NDD phenotype although their regulatory roles were not studied *in vitro*. Other Del donors doesn’t carry more than two enSNP (one GH16J029819 SNP in Del3, one GH16J030112 SNP in Del4). In the future, more hiPSC clones from 16p11.2del patients carrying same residual haplotype as Del1 will be collected and *in vitro* functional assay will be performed to confirm the cis-regulation of enSNP to other 16p11.2 genes.

## LIMITATIONS

First, hiPSC clones of 16p11.2del carriers in this study was limited, and more hiPSC clones of intra-family 16p11.2del carriers should be recruited to replicate our residual haplotype-specific cellular phenotypes and identify its modifier effect to other 16p11.2 gene. Second, the clonal variant between intra- and inter-donor has been reported (Brennand et al., 2011) and should be avoided or studied unbiased by single-cell RNA-Seq as reported from *NRXN1* deletion (Lam et al., 2019). Third, spatial DNA contact is important step for long-range regulation of gene expression during human brain (Won et al., 2016; de la Torre-Ubieta et al., 2018). The cis- and trans-acting chromatin contacts of 16p11.2del (BP2-BP3) (Loviglio et al., 2017) and 22q11.2del (Zhang et al., 2018) to adjacent genes or whole genome have been reported in lymphoblastoid cell lines (LCLs) of affected patient. Consequently, mapping the spatial regulation landscape of 16p11.2del in hiPS-derived neuron cell or organoid should be performed to decode the genome-wide regulatory effect of 16p11.2 region during early neurogenesis.

## MATERIALS AND METHODS

### Subjects

31 NDD patients with 16p11.2del from 29 independent families were recruited in CIP-NDD-16p11.2del cohort. 16p11.2del were detected by either CMA testing, MLPA or exome sequencing. The neurodevelopmental quotient (DQ) was evaluated using Wechsler Intelligence Scale for Children (WISC), or the Children Neuropsychological and Behavior Scale-Revision 2016 (CNBS-R2016) which is developed by the Capital Institute of Pediatrics to assess the developmental level of children aged 0-6 years and is widely used in China (Li et al., 2019). DSM-IV was used for ASD diagnosis. The procedure was reviewed and approved by the ethics committee of the Capital Institute of Pediatrics. All individuals or guardians have signed written informed consent for donating peripheral blood and the publication of related works.

### Target sequencing of 16p11.2 interval and phasing residual haplotype of 16p11.2 locus

The DNA was extracted from peripheral leukocytes and target regions were baited (244.393 kb, chr16: 29592784-30200783, hg19 version, the location of 2077 bait probes were in Supplementary File 1) using one commercial capture kit (MyGenostics GenCap Enrichment Technologies). The enriched libraries were sequenced on Illumina HiSeq X Ten sequencer (Illumina) for paired reads of 150 bp. Illumina sequencing adapters and low-quality reads were filtered out and the clean reads were mapped to the UCSC hg19 version and processed using GATK software (https://gatk.broadinstitute.org). The data were then transformed to variant call format (VCF). The average coverage of tested samples ranged from 162x to 2265x, and the percent of target areas >20x ranged from 88.94% to 97.22%.

With online annotation software (QIAGEN, Clinical Insight (QCI) Interpret Translational tool, https://apps.qiagenbioinformatics.cn/), the homozygous high-quality variants with MAF 10-90% were extracted to phase accurate residual haplotype for 16p11.2del carriers, combining the intra-family Mendeline inheritance model (Table S1). Besides public genomic databases (1000 Genomes, dbSNP, gnomAD), one commercial Chinese whole genome sequencing database (DeYi WGS database of 690 Chinese normal population) were used to annotate Chinese minor allele frequency (MAF) of 16p11.2 region. Beside common variant, rare SNVs (MAF<10%) were also extracted in order to exclude other deleterious or pathogenic coding variants (loss of function variant or missense variant with REVEL>0.7).

Ten 16p11.2del iPS donors of Julien G Roth et al. (Roth et al., 2020) studied were from Simons Foundation Autism Research Initiative (SFARI, https://www.sfari.org/). We applied SFARI Base (SFARI Request ID 12603.1.3) and were accessed for SNP-chip information of these donors and phased the residual haplotypes for 16p11.2del donors (16 SNPs were used to phase residual haplotype of SFARI samples).

### Generation of human induced pluripotent stem cells (hiPSCs)

PBMCs were reprogrammed to hiPSCs using Invitrogen CytoTune-iPS 2.0 Sendai reprogramming kit (Invitrogen), which contained OCT3/4, KLF4, SOX2 and cMYC as reprogramming factors and has been previously reported (Seki et al., 2010). In brief, PBMCs were seeded into 12-well plates and cultured for 14 days in PBMC medium. Then, 2.5×10^5^ PBMCs were infected with Sendai virus and cultured for 48h at 37°C. Further, infected cells were transferred to mouse embryonic fibroblast (MEF) feeders in iPSCs medium. After 2 weeks, individual iPSC colonies were picked manually and transferred to Matrigel (Corning)-coated maintained in mTeSR medium (Stem Cell Technologies). Cells were tested routinely to be mycoplasma-negative. Only the iPSCs more than passage 10 were characterized, including pluripotency analysis and genomic background. For pluripotency analysis, we performed AP staining, immunofluorescent staining and teratoma formation. For genomic background, we performed karyotype, short tandem repeat (STR) site analysis and array CGH to confirm identical genome. The relationship of two families were confirmed by STR.

### Differentiation of cortical neuron cells

iPSCs colonies were differentiated into neurons with modified protocol (Zhang et al., 2013). The time before Day0 was labeled as minus (see Fig. 2C). On Day-15, hiPSCs was seeded in an Matrigel-coated (Corning) 6-well plates in mTeSR medium supplemented with 10 μM Y-27632. Following 24h, if the cells reached ~60% confluence, hiPSCs were then infected with NGN2-expressing lentiviruses (10ml mTeSR+5ul polybrene +2.5μl lentiviruses) for 18h. Culture medium was replaced with fresh mTeSR medium for 24h. After this, 1mg/mL puromycin was added for 48h to select for successfully transduced cells. After 1-2 times of passages, iPSCs were treated with accutase and plated as dissociated cells in Matrigel-coated 6-well plates (Day-5). On Day-4, Doxycycline was added to induce NGN2 expression and retained in the medium until Day0, getting neural precursor cells (NPCs). On Day0, NPCs were dissociated into single cells with accutase and seeded onto 24-well plates (1×10^5^ cells) in NPCs medium (49ml Neurobasal medium+1ml B27), and the plates were coated in 0.07% PEI for 1h at room temperature, washed with sterile water, and dried overnight. Then the plates were coated with 10 μg/ml laminin at 4°C overnight. On Day1 (<24h after plating), the NPCs medium was removed and replaced with neuronal cells (NCs) medium (49ml Neurobasal medium+1ml B27+50ul GDNF (10μg/mL) + 50ul BDNF (10μg/mL) + 50ul cAMP(50mM) + 50ulVc (0.2mM)). After Day2, 50% of the NCs medium in each well was exchanged every 2-3 days. The validation of mature NCs was on Day21. Fig 2C presented the schematic of reprogramming of cortical neuron cells.

### RNA sequencing and transcriptomic analysis

RNA was collected ontDay0 and Day21 (red ovals in Fig. 2C) respectively using RNeasy Plus Mini Kit (QIAGEN) according to the manufacturer’s protocol with the step of on-column DNA digestion (QIAGEN) to remove genomic DNA. RNA integrity was verified on an Agilent 2100 Bioanalyzer, and the concentration was determined using Invitrogen Qubit 3.0 Spectrophotometer. Then RNA libraries were prepared using TruSeq RNA sample Preparation kit V2 (Illumina) and sequenced on an Illumina Hiseq 2500 to generate paired-end 150bp bases in length (Genesky Biotechnologies Inc, Shanghai). Initial sequencing data quality was assessed using Fast QC (v0.11.4). With Fastx-Toolkit (v0.0.13), further read trimming was performed to get clean reads. The trimmed reads were aligned to the reference genome hg19 using HISAT2 (v2.2.1). After that, RSeQC-2.3.2 was used to assess saturation analysis, duplicate reads and RNA degradation analysis. Lastly, raw read counts and FPKM for genes was extracted using String Tie (v2.1.2), which were used for the subsequent bioinformatic analysis. Two or three clones per each donor finished RNA-Seq. One normal sample’s PBMC was performed for RNA-Seq. All RNA-Seq data were deposited in Chinese National Genomics Data Center with accession number (https://ngdc.cncb.ac.cn/omix, HRA002368).

Principal component analysis (PCA) was used to evaluate the quality of transcriptome. Surrogate variable analysis (SVA) was performed to correct the effects of batch and additional uncharacterized factors, and all detected genes were included for PCA. Pheatmap (v1.0.10) was used to visualize the expression of 16p11.2 interval genes, the markers of NPCs and NCs. The values were transformed from FPKM values by zFPKM (v.1.16.0)

### Weighted gene co-expression network analysis (WGCNA) analysis

Weighted gene co-expression network analysis (WGCNA, WGCNA package v.1.70-3) in R (v.4.1.2) (Langfelder and Horvath, 2008) was performed to identify the pattern of significant genes and related clinical feature. Primarily, we analyzed the adjacency matrix, calculated the topological overlap matrix and performed hierarchical clustering. After that, modules were identified using the dynamic tree cut algorithm, and the first-principal component was calculated with the module eigengene (MEs) function. With the first-principal component, MEss were determined. Further, Module eigengene-based connectivity (kME) was determined by calculating the correlation between gene expression and the ME and the correlations between modules and traits was represented as Pearson Correlations. For the genes in the modules with the greatest correlations with the neurodevelopmental phenotype, enrichment analysis was finished using R package (clusterProfiler v.4.2.2).

Differential expression analysis in each pair-wise comparison was performed using DEseq2. Genes were identified as differentially expressed if the fold change was higher than 1.5 and Benjamini-Hochberg adjusted p-value fell below 0.05. For the differential expressed genes (DEGs), pathway enrichment analysis was performed using the R package (clusterProfiler v.4.2.2), including gene ontology and KEGG pathways.

### Multi-electrode array (MEA) recording and analysis

Multi-well MEA is high-throughput assay to reflect the electrophysiological properties of neural or heart cells (Nam and Wheeler, 2011). Spontaneous electrical activities of MEA plates were collected periodically. Four days after doxycycline induction of NGN2, 1×10^5^ NPCs (per well) were seeded onto previously 0.3% PEI solution+10μg/ml laminin-coated 24-well MEA plates (Axion Biosystems). On Day1 (dashed red rectangle in Fig. 2C), NPCs medium was replaced with NC medium, and, thereafter, half medium changes were conducted every 2-3 days. NPCs interfered with by lentivirus expressing *MAPK3*, *PRRT2* or blank were prepared and plated in the same manner. During MEA recording, plate was incubated for 5 minutes on the 37°C pre-warmed reader with half-refreshed NCs media. The bandpass filter was 200 Hz-3 kHz. The recording data were analyzed using Axis Navigator1.5 software. The weighted mean firing rates was compared by Analysis of Variance of Aligned Rank Transformed Data (Wobbrock, 2011), and p<0.05 was considered as statistically significant.

### Immunofluorescence experiments

Cultured cells were fixed with 4% paraformaldehyde (PFA) for 20 min, incubated in 0.5% Triton X-100 in PBS for 30min and blocked within goat serum for 1h. Then cells were incubated with the primary antibody solution (Table S19) overnight at 4°C and fluorophore-conjugated secondary antibodies for 1h at room temperature. Lastly, nuclei were counterstained with DAPI or Hoechst 33342. Images were taken using Nikon or Leica confocal microscope. Images were analyzed with Image J/Fiji software.

### Reverse-Transcription (RT) and qPCR

Reverse-Transcription (RT) following qPCR or PCR was performed to validate the expression of 16p11.2 genes (*PRRT2, TLCD3B, PAGR1, SEZ6L2, CDIPT, MAPK3, YPEL3, TAOK2, KCTD13* and *MVP*) and potential enhancer RNAs (eRNAs) of 16p11.2 region. RNA was isolated as described above following genomic DNA digestion. Purified total RNA (1000ng/sample) was used as the template for RT reaction, and the diluted (1:100) RT product was used as template for qPCR reaction. The qPCR was performed on a 7500 FAST Real-Time PCR system (Life Technologies) using SYBR® Select Master Mix kit (Life Technologies) with specific primers. Fold changes were calculated using the Ct method, and the average expression was compared in each pair-wise comparison. The RT-qPCR primers were listed in Table S17.

Eight enhancer RNAs (eRNAs) were chosen for reverse-transcription PCR (RT-PCR, eRNAs products in Fig. 8A, primers and sizes of designed PCR products in Table S17). Sanger sequencing and alignment of amplicons to were used to confirm the genomic locations of these eRNAs.

### Western blot

Cultured NPCs cells were lysed to RIPA-lysis buffer (Santa Cruz Biotechnology) supplemented with protease and/or phosphatase inhibitor (Thermo scientific). Lysates were centrifuged (4°C, 13000rpm,10 min) to collect protein and the concentration of protein was determined with the BCA assay (Thermo Fisher Scientific). Western blot was performed following the manufacture’s instruction of the corresponding antibodies (Table S19). Besides, NPCs with 16p11.2del interfered with by lentiviruses expressing *MAPK3*, *PRRT2* or blank were tested in the same manner.

Quantification of Western blots was performed using ImageJ/Fiji software. Statistical analysis in each pair-wise comparison was determined using unpaired t-tests, and the two-sided p value less than 0.05 was considered as statistically significant.

### Rescue assays

The rescue of 16p11.2 NPCs was achieved by designed lentivirus-mediated *MAPK3, PRRT2* or blank overexpression systems (Cherry/GFP-*PRRT2/MAPK3/blank*) (Genechem), with titer of 1×10^8^ TU/ml (*MAPK3*) or 1.2×0^8^ TU/ml (*PRRT2*). We performed lentiviral transfection at Day-3 (red rectangle in Fig. 2B) with two different doses of lentivirus (dose of 2×10^4^ TU for *MAPK3* or 2.4×10^4^ TU for *PRRT2* was abbreviated as R (+); dose of 1×10^5^ TU for *MAPK3* or 1.2×10^5^ TU for *PRRT2* was abbreviated as R (++)). After 48hr transfection, the NPCs were lysed for western blotting or plated on MEA recording electrophysiological properties. During the differentiation process of cortical neuron cell as described above, Doxycycline was added to induce NGN2 expression on Day4. After 24h, medium was removed and cells were infected by the addition of lentiviral vectors (0.2 or 1ul) into N2 medium for 18h. The infected cells were cultured in medium supplemented with Doxycycline for 48h to get NPCs, which were used to the subsequent experiments, including MEA recording and western blot. Given that firing spikes will be recorded till Day33, we plated NPCs with R (++) dosage during MEA experiment and collected firing signals of NCs from Day9 to Day33 for comparison.

### Vector construction and luciferase reporter assay

Among ten SNPs covering elite regulatory elements, eight enhancer SNPs (GH16J029925 and GH16J029972) were chosen for *in vitro* luciferase reporter assay because multiple SNPs on same enhancer. DNA of Del1 and Del2 from Family 1 were used to amplify different haplotypic fragments with long-range PCR or TD-PCR (Table S17) adding restriction enzymes (MluI and KpnI, NEB) on the 3’ of primers. We constructed long fragment covering entire GH16J029925 fragment (chr16:29936436-29938999, 2495bp) and four enhancer SNPs (rs12716972/rs1302842915/rs12716973/rs4420550). GH16J029972 is too large to be constructed, we compromised to construct one long fragment (chr16:29983649-29985275, 1627bp, rs11901/rs4318227) and two short fragments (chr16: 29986384-29986709, 326bp; chr16:29988775-29989141, 367bp) covering rs11150577 and rs4583255 respectively. Only one 1627bp regulatory fragment was failed in amplification due to repetitive sequences (black box in Fig. 8A, labeled yellow in Table S17). After sequencing validation, these regulatory fragments were cloned to the pGL3-promoter vector. The spontaneous mutation during the PCR due to high GC content was excluded or artificial correction.

*In vitro* luciferase reporter assay was conducted in the HEK293T cells (human embryonic kidneys 293T) to test the effects of different residual haplotype on DNA enhancer activities. Twenty-four hours post-transfection using Lipofectamine 3000 (Thermo Scientific), transfected cells were collected, lysed. The luciferase activity was determined by Dual-Luciferase Reporter Assay System (Promega). The firefly luciferase activity was normalized to that of Renilla luciferase to control for transfection efficiency variations.

### Residual haplotype phasing for both ASD and CS patients with 16p11.2del

We applied SFARI Base and were accessed (SFARI Request ID 12603.1.3) for the phenotypic and genotypic information of 86 carriers with 16p11.2del (SFARI-16p11.2del) from Phase 1 Dataset (SFARI Base, https://www.sfari.org/), including 18 ASD patients (SFARI-ASD-16p11.2del cohort). 16 SNPs of test haplotype were available for genotype phasing, and they showed closely linked for 72.1% carriers (62/86 in SFARI-16p11.2del and 13/18 in SFARI-ASD-16p11.2del).

29 CS patients with 16p11.2del were diagnosed and recruited to the PUMC-CS-16p11.2del cohort in Department of Orthopedic Surgery, Peking Union Medical College Hospital. All patients have been performed whole genome sequencing which are available to phase residual haplotype. 58 SNPs of test haplotype showed closely linked for 89.7% patients (26/29).

Ten 16p11.2del iPS donors of Julien G Roth et al. (Roth et al., 2020) were also from Simons Foundation Autism Research Initiative (SFARI, https://www.sfari.org/). Their residual haplotypes were phased as above.

## Availability of data and materials

The RNA-Seq datasets used for the study were deposited in Chinese National Genomics Data Center (https://ngdc.cncb.ac.cn/omix, HRA002368) and are available from corresponding author upon request.

## Distribution of materials

hiPSCs is stored on China Center for Type Culture Collection (CCTCC, http://www.cctcc.org) and their distribution are under permission from the ethics committee of the Capital Institute of Pediatrics.

## Acknowledgements

We appreciated obtaining access to [Simons Searchlight 16p11.2 Phase 2 and Phase 1 Dataset] data on SFARI Base (https://www.sfari.org/), and appreciated Chigene (Beijing) Translational Medical Research Center Co., Ltd for 600 Chinese WGS data. This work was supported by grants from the Beijing Natural Science Foundation (7202019 to Xiaoli Chen), the Chinese National Nature Science Fund (31671310 to Xiaoli Chen); Capital Health Research and Development of Special (2020-1-4071 and 2020-2-1131), the Innovation Project of Beijing Municipal Human Resources and Social Security Bureau to Xiaoli Chen. Research Foundation of Capital Institute of Pediatrics (CXYJ-2021006 to Xiaoli Chen).

## Declaration

The authors declare no conflicts of interest.

## References

Adamo, A., Atashpaz, S., Germain, P.L., Zanella, M., D’Agostino, G., Albertin, V., et al. (2015). 7q11.23 dosage-dependent dysregulation in human pluripotent stem cells affects transcriptional programs in disease-relevant lineages. Nat Genet 47(2), 132–141. doi: 10.1038/ng.3169.

Andersson, R., Gebhard, C., Miguel-Escalada, I., Hoof, I., Bornholdt, J., Boyd, M., et al. (2014). An atlas of active enhancers across human cell types and tissues. Nature 507(7493), 455–461. doi: 10.1038/nature12787.

Arbogast, T., Ouagazzal, A.M., Chevalier, C., Kopanitsa, M., Afinowi, N., Migliavacca, E., et al. (2016). Reciprocal Effects on Neurocognitive and Metabolic Phenotypes in Mouse Models of 16p11.2 Deletion and Duplication Syndromes. PLoS Genet 12(2), e1005709. doi: 10.1371/journal.pgen.1005709.

Arbogast, T., Razaz, P., Ellegood, J., McKinstry, S.U., Erdin, S., Currall, B., et al. (2019). Kctd13-deficient mice display short-term memory impairment and sex-dependent genetic interactions. Hum Mol Genet 28(9), 1474–1486. doi: 10.1093/hmg/ddy436.

Barrett, J.C., Fry, B., Maller, J., and Daly, M.J. (2005). Haploview: analysis and visualization of LD and haplotype maps. Bioinformatics 21(2), 263–265. doi: 10.1093/bioinformatics/bth457.

Blizinsky, K.D., Diaz-Castro, B., Forrest, M.P., Schurmann, B., Bach, A.P., Martin-de-Saavedra, M.D., et al. (2016). Reversal of dendritic phenotypes in 16p11.2 microduplication mouse model neurons by pharmacological targeting of a network hub. Proc Natl Acad Sci U S A 113(30), 8520–8525. doi: 10.1073/pnas.1607014113.

Blumenthal, I., Ragavendran, A., Erdin, S., Klei, L., Sugathan, A., Guide, J.R., et al. (2014). Transcriptional consequences of 16p11.2 deletion and duplication in mouse cortex and multiplex autism families. Am J Hum Genet 94(6), 870–883. doi: 10.1016/j.ajhg.2014.05.004.

Brennand, K.J., Simone, A., Jou, J., Gelboin-Burkhart, C., Tran, N., Sangar, S., et al. (2011). Modelling schizophrenia using human induced pluripotent stem cells. Nature 473(7346), 221–225. doi: 10.1038/nature09915.

Coe, B.P., Stessman, H.A.F., Sulovari, A., Geisheker, M.R., Bakken, T.E., Lake, A.M., et al. (2019). Neurodevelopmental disease genes implicated by de novo mutation and copy number variation morbidity. Nat Genet 51(1), 106–116. doi: 10.1038/s41588-018-0288-4.

Cooper, G.M., Coe, B.P., Girirajan, S., Rosenfeld, J.A., Vu, T.H., Baker, C., et al. (2011). A copy number variation morbidity map of developmental delay. Nat Genet 43(9), 838–846. doi: 10.1038/ng.909.

de la Torre-Ubieta, L., Stein, J.L., Won, H., Opland, C.K., Liang, D., Lu, D., et al. (2018). The Dynamic Landscape of Open Chromatin during Human Cortical Neurogenesis. Cell 172(1-2), 289–304 e218. doi: 10.1016/j.cell.2017.12.014.

Deshpande, A., Yadav, S., Dao, D.Q., Wu, Z.Y., Hokanson, K.C., Cahill, M.K., et al. (2017). Cellular Phenotypes in Human iPSC-Derived Neurons from a Genetic Model of Autism Spectrum Disorder. Cell Rep 21(10), 2678–2687. doi: 10.1016/j.celrep.2017.11.037.

Drakulic, D., Djurovic, S., Syed, Y.A., Trattaro, S., Caporale, N., Falk, A., et al. (2020). Copy number variants (CNVs): a powerful tool for iPSC-based modelling of ASD. Mol Autism 11(1), 42. doi: 10.1186/s13229-020-00343-4.

Duyzend, M.H., Nuttle, X., Coe, B.P., Baker, C., Nickerson, D.A., Bernier, R., et al. (2016). Maternal Modifiers and Parent-of-Origin Bias of the Autism-Associated 16p11.2 CNV. Am J Hum Genet 98(1), 45–57. doi: 10.1016/j.ajhg.2015.11.017.

Egolf, L.E., Vaksman, Z., Lopez, G., Rokita, J.L., Modi, A., Basta, P.V., et al. (2019). Germline 16p11.2 Microdeletion Predisposes to Neuroblastoma. Am J Hum Genet 105(3), 658–668. doi: 10.1016/j.ajhg.2019.07.020.

Escamilla, C.O., Filonova, I., Walker, A.K., Xuan, Z.X., Holehonnur, R., Espinosa, F., et al. (2017). Kctd13 deletion reduces synaptic transmission via increased RhoA. Nature 551(7679), 227–231. doi: 10.1038/nature24470.

Fishilevich, S., Nudel, R., Rappaport, N., Hadar, R., Plaschkes, I., Iny Stein, T., et al. (2017). GeneHancer: genome-wide integration of enhancers and target genes in GeneCards. Database (Oxford) 2017. doi: 10.1093/database/bax028.

Frega, M., Linda, K., Keller, J.M., Gumus-Akay, G., Mossink, B., van Rhijn, J.R., et al. (2019). Neuronal network dysfunction in a model for Kleefstra syndrome mediated by enhanced NMDAR signaling. Nat Commun 10(1), 4928. doi: 10.1038/s41467-019-12947-3.

Giannuzzi, G., Schmidt, P.J., Porcu, E., Willemin, G., Munson, K.M., Nuttle, X., et al. (2019). The Human-Specific BOLA2 Duplication Modifies Iron Homeostasis and Anemia Predisposition in Chromosome 16p11.2 Autism Individuals. Am J Hum Genet 105(5), 947–958. doi: 10.1016/j.ajhg.2019.09.023.

Gibson, G., Powell, J.E., and Marigorta, U.M. (2015). Expression quantitative trait locus analysis for translational medicine. Genome Med 7(1), 60. doi: 10.1186/s13073-015-0186-7.

Girdhar, K., Hoffman, G.E., Jiang, Y., Brown, L., Kundakovic, M., Hauberg, M.E., et al. (2018). Cell-specific histone modification maps in the human frontal lobe link schizophrenia risk to the neuronal epigenome. Nat Neurosci 21(8), 1126–1136. doi: 10.1038/s41593-018-0187-0.

Girirajan, S., Rosenfeld, J.A., Coe, B.P., Parikh, S., Friedman, N., Goldstein, A., et al. (2012). Phenotypic heterogeneity of genomic disorders and rare copy-number variants. N Engl J Med 367(14), 1321–1331. doi: 10.1056/NEJMoa1200395.

Golzio, C., Willer, J., Talkowski, M.E., Oh, E.C., Taniguchi, Y., Jacquemont, S., et al. (2012). KCTD13 is a major driver of mirrored neuroanatomical phenotypes of the 16p11.2 copy number variant. Nature 485(7398), 363–367. doi: 10.1038/nature11091.

Harel, T., and Lupski, J.R. (2018). Genomic disorders 20 years on-mechanisms for clinical manifestations. Clin Genet 93(3), 439–449. doi: 10.1111/cge.13146.

Hudac, C.M., Bove, J., Barber, S., Duyzend, M., Wallace, A., Martin, C.L., et al. (2020). Evaluating heterogeneity in ASD symptomatology, cognitive ability, and adaptive functioning among 16p11.2 CNV carriers. Autism Res 13(8), 1300–1310. doi: 10.1002/aur.2332.

Iyer, J., Singh, M.D., Jensen, M., Patel, P., Pizzo, L., Huber, E., et al. (2018). Pervasive genetic interactions modulate neurodevelopmental defects of the autism-associated 16p11.2 deletion in Drosophila melanogaster. Nat Commun 9(1), 2548. doi: 10.1038/s41467-018-04882-6.

Jacquemont, S., Reymond, A., Zufferey, F., Harewood, L., Walters, R.G., Kutalik, Z., et al. (2011). Mirror extreme BMI phenotypes associated with gene dosage at the chromosome 16p11.2 locus. Nature 478(7367), 97–102. doi: 10.1038/nature10406.

Lam, M., Moslem, M., Bryois, J., Pronk, R.J., Uhlin, E., Ellstrom, I.D., et al. (2019). Single cell analysis of autism patient with bi-allelic NRXN1-alpha deletion reveals skewed fate choice in neural progenitors and impaired neuronal functionality. Exp Cell Res 383(1), 111469. doi: 10.1016/j.yexcr.2019.06.014.

Langfelder, P., and Horvath, S. (2008). WGCNA: an R package for weighted correlation network analysis. BMC Bioinformatics 9, 559. doi: 10.1186/1471-2105-9-559.

Lee, H.Y., Huang, Y., Bruneau, N., Roll, P., Roberson, E.D., Hermann, M., et al. (2012). Mutations in the gene PRRT2 cause paroxysmal kinesigenic dyskinesia with infantile convulsions. Cell Rep 1(1), 2–12. doi: 10.1016/j.celrep.2011.11.001.

Levo, M., and Segal, E. (2014). In pursuit of design principles of regulatory sequences. Nat Rev Genet 15(7), 453–468. doi: 10.1038/nrg3684.

Li, H.H., Feng, J.Y., Wang, B., Zhang, Y., Wang, C.X., and Jia, F.Y. (2019). Comparison Of The Children Neuropsychological And Behavior Scale And The Griffiths Mental Development Scales When Assessing The Development Of Children With Autism. Psychol Res Behav Manag 12, 973–981. doi: 10.2147/PRBM.S225904.

Liu, Y.T., Nian, F.S., Chou, W.J., Tai, C.Y., Kwan, S.Y., Chen, C., et al. (2016). PRRT2 mutations lead to neuronal dysfunction and neurodevelopmental defects. Oncotarget 7(26), 39184–39196. doi: 10.18632/oncotarget.9258.

Loviglio, M.N., Leleu, M., Mannik, K., Passeggeri, M., Giannuzzi, G., van der Werf, I., et al. (2017). Chromosomal contacts connect loci associated with autism, BMI and head circumference phenotypes. Mol Psychiatry 22(6), 836–849. doi: 10.1038/mp.2016.84.

Moreno-De-Luca, A., Evans, D.W., Boomer, K.B., Hanson, E., Bernier, R., Goin-Kochel, R.P., et al. (2015). The role of parental cognitive, behavioral, and motor profiles in clinical variability in individuals with chromosome 16p11.2 deletions. JAMA Psychiatry 72(2), 119–126. doi: 10.1001/jamapsychiatry.2014.2147.

Nam, Y., and Wheeler, B.C. (2011). In vitro microelectrode array technology and neural recordings. Crit Rev Biomed Eng 39(1), 45–61. doi: 10.1615/critrevbiomedeng.v39.i1.40.

Niarchou, M., Chawner, S., Doherty, J.L., Maillard, A.M., Jacquemont, S., Chung, W.K., et al. (2019). Psychiatric disorders in children with 16p11.2 deletion and duplication. Transl Psychiatry 9(1), 8. doi: 10.1038/s41398-018-0339-8.

Park, S.M., Park, H.R., and Lee, J.H. (2017). MAPK3 at the Autism-Linked Human 16p11.2 Locus Influences Precise Synaptic Target Selection at Drosophila Larval Neuromuscular Junctions. Mol Cells 40(2), 151–161. doi: 10.14348/molcells.2017.2307.

Portmann, T., Yang, M., Mao, R., Panagiotakos, G., Ellegood, J., Dolen, G., et al. (2014). Behavioral abnormalities and circuit defects in the basal ganglia of a mouse model of 16p11.2 deletion syndrome. Cell Rep 7(4), 1077–1092. doi: 10.1016/j.celrep.2014.03.036.

Pucilowska, J., Vithayathil, J., Pagani, M., Kelly, C., Karlo, J.C., Robol, C., et al. (2018). Pharmacological Inhibition of ERK Signaling Rescues Pathophysiology and Behavioral Phenotype Associated with 16p11.2 Chromosomal Deletion in Mice. J Neurosci 38(30), 6640–6652. doi: 10.1523/JNEUROSCI.0515-17.2018.

Pucilowska, J., Vithayathil, J., Tavares, E.J., Kelly, C., Karlo, J.C., and Landreth, G.E. (2015). The 16p11.2 deletion mouse model of autism exhibits altered cortical progenitor proliferation and brain cytoarchitecture linked to the ERK MAPK pathway. J Neurosci 35(7), 3190–3200. doi: 10.1523/JNEUROSCI.4864-13.2015.

Rein, B., and Yan, Z. (2020). 16p11.2 Copy Number Variations and Neurodevelopmental Disorders. Trends Neurosci 43(11), 886–901. doi: 10.1016/j.tins.2020.09.001.

Richter, M., Murtaza, N., Scharrenberg, R., White, S.H., Johanns, O., Walker, S., et al. (2019). Altered TAOK2 activity causes autism-related neurodevelopmental and cognitive abnormalities through RhoA signaling. Mol Psychiatry 24(9), 1329–1350. doi: 10.1038/s41380-018-0025-5.

Roth, J.G., Muench, K.L., Asokan, A., Mallett, V.M., Gai, H., Verma, Y., et al. (2020). 16p11.2 microdeletion imparts transcriptional alterations in human iPSC-derived models of early neural development. Elife 9. doi: 10.7554/eLife.58178.

Seki, T., Yuasa, S., Oda, M., Egashira, T., Yae, K., Kusumoto, D., et al. (2010). Generation of induced pluripotent stem cells from human terminally differentiated circulating T cells. Cell Stem Cell 7(1), 11–14. doi: 10.1016/j.stem.2010.06.003.

Shcheglovitov, A., Shcheglovitova, O., Yazawa, M., Portmann, T., Shu, R., Sebastiano, V., et al. (2013). SHANK3 and IGF1 restore synaptic deficits in neurons from 22q13 deletion syndrome patients. Nature 503(7475), 267–271. doi: 10.1038/nature12618.

Shen, Y., Chen, X., Wang, L., Guo, J., Shen, J., An, Y., et al. (2011). Intra-family phenotypic heterogeneity of 16p11.2 deletion carriers in a three-generation Chinese family. Am J Med Genet B Neuropsychiatr Genet 156(2), 225–232. doi: 10.1002/ajmg.b.31147.

Shinawi, M., Liu, P., Kang, S.H., Shen, J., Belmont, J.W., Scott, D.A., et al. (2010). Recurrent reciprocal 16p11.2 rearrangements associated with global developmental delay, behavioural problems, dysmorphism, epilepsy, and abnormal head size. J Med Genet 47(5), 332–341. doi: 10.1136/jmg.2009.073015.

Simons Vip, C. (2012). Simons Variation in Individuals Project (Simons VIP): a genetics-first approach to studying autism spectrum and related neurodevelopmental disorders. Neuron 73(6), 1063–1067. doi: 10.1016/j.neuron.2012.02.014.

Song, M., Yang, X., Ren, X., Maliskova, L., Li, B., Jones, I.R., et al. (2019). Mapping cis-regulatory chromatin contacts in neural cells links neuropsychiatric disorder risk variants to target genes. Nat Genet 51(8), 1252–1262. doi: 10.1038/s41588-019-0472-1.

Steinberg, S., de Jong, S., Mattheisen, M., Costas, J., Demontis, D., Jamain, S., et al. (2014). Common variant at 16p11.2 conferring risk of psychosis. Mol Psychiatry 19(1), 108–114. doi: 10.1038/mp.2012.157.

Sundberg, M., Pinson, H., Smith, R.S., Winden, K.D., Venugopal, P., Tai, D.J.C., et al. (2021). 16p11.2 deletion is associated with hyperactivation of human iPSC-derived dopaminergic neuron networks and is rescued by RHOA inhibition in vitro. Nat Commun 12(1), 2897. doi: 10.1038/s41467-021-23113-z.

Tai, D.J., Ragavendran, A., Manavalan, P., Stortchevoi, A., Seabra, C.M., Erdin, S., et al. (2016). Engineering microdeletions and microduplications by targeting segmental duplications with CRISPR. Nat Neurosci 19(3), 517–522. doi: 10.1038/nn.4235.

Thomas, G.M., and Huganir, R.L. (2004). MAPK cascade signalling and synaptic plasticity. Nat Rev Neurosci 5(3), 173–183. doi: 10.1038/nrn1346.

Urresti, J., Zhang, P., Moran-Losada, P., Yu, N.K., Negraes, P.D., Trujillo, C.A., et al. (2021). Cortical organoids model early brain development disrupted by 16p11.2 copy number variants in autism. Mol Psychiatry. doi: 10.1038/s41380-021-01243-6.

Valente, P., Castroflorio, E., Rossi, P., Fadda, M., Sterlini, B., Cervigni, R.I., et al. (2016). PRRT2 Is a Key Component of the Ca(2+)-Dependent Neurotransmitter Release Machinery. Cell Rep 15(1), 117–131. doi: 10.1016/j.celrep.2016.03.005.

Verbitsky, M., Westland, R., Perez, A., Kiryluk, K., Liu, Q., Krithivasan, P., et al. (2019). The copy number variation landscape of congenital anomalies of the kidney and urinary tract. Nat Genet 51(1), 117–127. doi: 10.1038/s41588-018-0281-y.

Weiss, L.A., Shen, Y., Korn, J.M., Arking, D.E., Miller, D.T., Fossdal, R., et al. (2008). Association between microdeletion and microduplication at 16p11.2 and autism. N Engl J Med 358(7), 667–675. doi: 10.1056/NEJMoa075974.

Wobbrock, J.O., Findlater, L., Gergle, D., and Higgins, J. J. (2011). The aligned rank transform for nonparametric factorial analyses using only ANOVA procedures. Proceedings of the ACM Conference on Human Factors in Computing Systems (CHI ’11). Vancouver, British Columbia (May 7–12, 2011), 143–146. doi: 10.1145/1978942.1978963.

Won, H., de la Torre-Ubieta, L., Stein, J.L., Parikshak, N.N., Huang, J., Opland, C.K., et al. (2016). Chromosome conformation elucidates regulatory relationships in developing human brain. Nature 538(7626), 523–527. doi: 10.1038/nature19847.

Wu, N., Ming, X., Xiao, J., Wu, Z., Chen, X., Shinawi, M., et al. (2015). TBX6 null variants and a common hypomorphic allele in congenital scoliosis. N Engl J Med 372(4), 341–350. doi: 10.1056/NEJMoa1406829.

Xie, H., Liu, F., Zhang, Y., Chen, Q., Shangguan, S., Gao, Z., et al. (2020). Neurodevelopmental trajectory and modifiers of 16p11.2 microdeletion: A follow-up study of four Chinese children carriers. Mol Genet Genomic Med 8(11), e1485. doi: 10.1002/mgg3.1485.

Yang, N., Wu, N., Dong, S., Zhang, L., Zhao, Y., Chen, W., et al. (2020). Human and mouse studies establish TBX6 in Mendelian CAKUT and as a potential driver of kidney defects associated with the 16p11.2 microdeletion syndrome. Kidney Int 98(4), 1020–1030. doi: 10.1016/j.kint.2020.04.045.

Yuan, H., Shangguan, S., Li, Z., Luo, J., Su, J., Yao, R., et al. (2021). CNV profiles of Chinese pediatric patients with developmental disorders. Genet Med 23(4), 669–678. doi: 10.1038/s41436-020-01048-y.

Zhang, X., Zhang, Y., Zhu, X., Purmann, C., Haney, M.S., Ward, T., et al. (2018). Local and global chromatin interactions are altered by large genomic deletions associated with human brain development. Nat Commun 9(1), 5356. doi: 10.1038/s41467-018-07766-x.

Zhang, Y., Pak, C., Han, Y., Ahlenius, H., Zhang, Z., Chanda, S., et al. (2013). Rapid single-step induction of functional neurons from human pluripotent stem cells. Neuron 78(5), 785–798. doi: 10.1016/j.neuron.2013.05.029.

Zufferey, F., Sherr, E.H., Beckmann, N.D., Hanson, E., Maillard, A.M., Hippolyte, L., et al. (2012). A 600 kb deletion syndrome at 16p11.2 leads to energy imbalance and neuropsychiatric disorders. J Med Genet 49(10), 660–668. doi: 10.1136/jmedgenet-2012-101203.

